# Structural basis for the ligand promiscuity of the neofunctionalized, carotenoid-binding fasciclin domain protein AstaP

**DOI:** 10.1101/2022.12.26.521925

**Authors:** Fedor D. Kornilov, Yury B. Slonimskiy, Daria A. Lunegova, Nikita A. Egorkin, Anna G. Savitskaya, Sergey Yu. Kleymenov, Eugene G. Maksimov, Sergey A. Goncharuk, Konstantin S. Mineev, Nikolai N. Sluchanko

**Author notes:** equal contributions (Fedor D. Kornilov, Yury B. Slonimskiy, Daria A. Lunegova). corresponding authors: Konstantin S. Mineev, Nikolai N. Sluchanko.

## Abstract

Fasciclins (FAS1) are ancient adhesion protein domains found across different phyla from bacteria to humans, with no common small ligand binding function reported. A unique FAS1-containing astaxanthin-binding protein (AstaP) from green algae can efficiently bind an unusually broad repertoire of carotenoids (astaxanthin, zeaxanthin, canthaxanthin, β-carotene), but the underlying mechanism is largely unknown. Here we dissect the structural basis for the ligand binding promiscuity of AstaP-orange1 (AstaPo1) by determining its solution NMR structure in complex with its natural ligand, astaxanthin (AXT), and validate this structure by SAXS, calorimetry, optical spectroscopy and mutagenesis data. While the unstructured tails of AstaPo1 are not essential for carotenoid binding, they enhance protein solubility. The a1-a2 helices of the AstaPo1 FAS1 domain embrace the carotenoid polyene like a jaw, organizing a conserved hydrophobic tunnel, too short to prevent the AXT β-ionone rings from protruding on both sides of the tunnel, thereby not imposing specificity restrictions. The only specific protein-AXT interactions involve H-bonds between the oxygenated groups on AXT and a peripheral Gln56 residue. Remarkably, mapping of this and other AXT-contacting AstaPo1 residues revealed their different conservation in AstaP orthologs with the tentative carotenoid-binding function and in FAS1 proteins in general, supporting neofunctionalization of AstaPs within green algae. Correspondingly, a cyanobacterial homolog with a similar domain structure cannot bind carotenoids due to subtle differences in residues decorating the tunnel. These structure-activity relationships inform the sequence-based prediction of the carotenoid-binding FAS1 members.

**SIGNIFICANCE:** A water-soluble astaxanthin-binding protein (AstaP) is a photoprotective protein in green algae helping them to tolerate stress conditions. While belonging to a ubiquitous protein family sharing an ancient structural domain, fasciclin, involved in cell adhesion, AstaP possesses an outstanding ability to bind carotenoid pigments of a different type, which are potent antioxidants. To understand the molecular basis for such carotenoid-binding promiscuity of AstaP, here we determined its spatial structure – the first structure of a carotenoid-protein complex solved by nuclear magnetic resonance spectroscopy. Together with biochemical and sequence conservation analyses, our data illustrate a remarkable case of neofunctionalization of the ancient protein domain and pave the way for its bioengineering and practical use as antioxidant transporter for biomedical applications.

## 1. Introduction

The fasciclin (FAS1) family comprises many proteins containing the FAS1-like domain (InterPro accession number IPR000782), which is heavily used in extracellular proteins serving as cell adhesion molecules (CAMs). The fasciclin I was first identified in insect CAMs within axon fascicles (hence the name fasciclin) (1) and can mediate homophilic or heterophilic cell adhesion, depending on whether FAS1 interacts with other FAS1 domains or with different CAMs such as integrins on the surface of other cells or with components of the extracellular matrix (2).

FAS1 domain is thought to have appeared in evolution very early and is ubiquitously found from bacteria and yeast to plants and animals (2). Such domains are typically 140 residues long and have a rather conserved α/β-mixed architecture belonging to a “β-grasp fold” superfamily (2). They are common in many secreted and transmembrane proteins and can be duplicated in one polypeptide. For example, four copies are found in insect fasciclin I (3), human periostin and transforming growth factor-β induced protein (2), and seven in human stabilin-1 and stabilin-2 (4)). Some proteins contain only a single FAS1 domain (for example, a mycobacterial immunogenic secreted protein MPB70 (5), a *Rhodobacter spheroides* protein Fdp (6) or a CupS protein from *Thermosynechococcus elongatus* (7)). Prokaryotes tend to have single-domain FAS1 proteins, whereas eukaryotes typically have such domains organized in tandems. Many FAS1-containing proteins are attached to the membrane and are glycosylated (2).

The first structure of a FAS1 domain was obtained for *Drosophila melanogaster* FAS1 (3). The fold of different FAS1 domains is common despite a rather low sequence identity among the members. All members of the FAS1 superfamily are characterized by the presence of conserved separate 10-15-residue motifs, H1 and H2, as well as a YH dyad in between (2). Although the predominant number of FAS1 structures was determined for isolated domains, there are seldom structures of multidomain FAS1 proteins such as that of the four-domain human transforming growth factor-β induced protein, mutations in which lead to abnormal self-association and accumulation of deposits in the cornea leading to dystrophy and blindness (8), or periostin, an extracellular matrix protein secreted by fibroblasts and up-regulated in a range of cancers (9). Nevertheless, the detailed molecular mechanism of the homophilic protein-protein recognition in cell adhesion involving FAS1 domains remains elusive.

That FAS1 domains are common in animals and plants became clear when the two-FAS1 domain CAMs homologous to *Drosophila* fasciclin I were identified in multicellular alga *Volvox carteri* (10). Higher plants were also shown to contain two-FAS1 domain CAMs (11). Interestingly, the cell adhesion function of FAS1 domains is so conserved that replacement of the original FAS1 domain by its distant homologs from either single-domain or multidomain FAS1 proteins does not abolish cell adhesion (12). At the same time, no common ligand binding function of the FAS1 domains has been reported.

In 2013, a remarkable protein was identified in green algae (13). While it was found as a stress-inducible protein containing a predicted FAS1 domain, it was isolated from the natural source as a soluble astaxanthin (AXT)-binding protein (hence called AstaP) undergoing glycosylation (13). While the preprotein contained a predicted signal peptide for secretion, it was accumulated on the cell periphery in response to hyperinsolation and osmotic stress, to perform a photoprotective role (13). Later on, several orthologs of AstaP have been isolated, which had either one or two FAS1 domains, differed by the pI and the presence of glycosylation sites (14, 15). Due to the differences in the absorbance spectrum, AstaP orthologs were classified into orange or pink subgroups (14). The recently described rather broad distribution of the new FAS1 family member, AstaP, in many *Scenedesmaceae* species provided compelling evidence toward its biologically relevant carotenoid-binding function (15). Yet, the carotenoid specificity had remained unaddressed.

AstaP was isolated from algal cells mainly as an AXT-bound form (16). However, focusing on the first AstaP described (AstaP-orange 1, or AstaPo1 for short) (13, 14), we have recently demonstrated that the recombinant AstaPo1 can bind a uniquely broad repertoire of carotenoids, which includes not only xanthophylls such as canthaxanthin (CAN) and zeaxanthin (ZEA), but also non-oxygenated β-carotene (βCar) (17) (**Fig. 1A**). AstaPo1 holoforms could be efficiently reconstituted *in vitro* by mixing with pure carotenoids or by expression in special carotenoid-synthesizing *E. coli* cells and could deliver the bound carotenoid to the apoforms of unrelated carotenoproteins or to biological membrane models (17). While these results expanded the toolkit of known carotenoproteins for carotenoid solubilization and targeted delivery, the structural basis for the carotenoid binding mechanism by AstaP, and potentially other FAS1 proteins, remained enigmatic. The apparent lack of carotenoid specificity also remained puzzling.

**Fig. 1.**
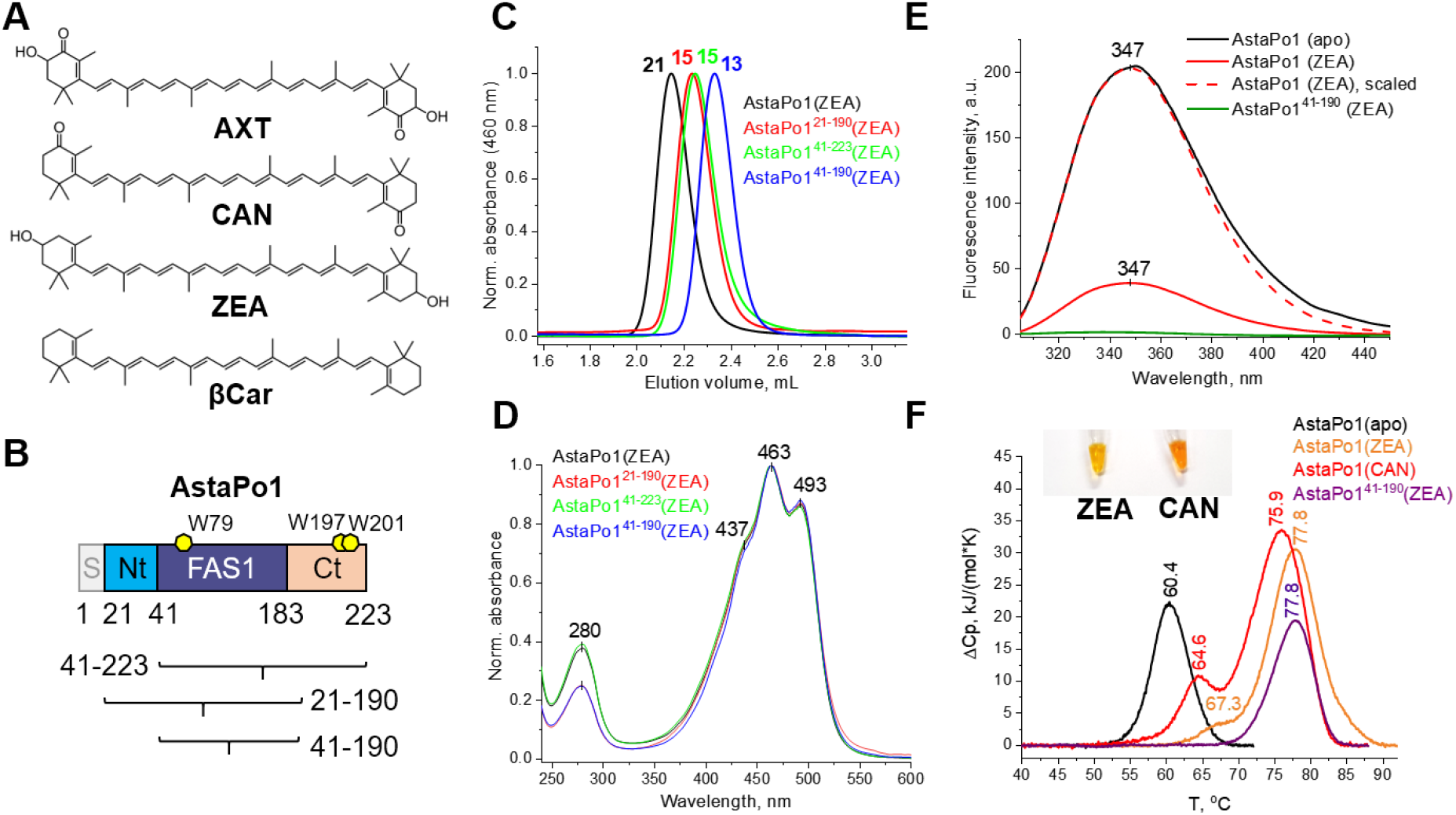
Localization of the carotenoid-binding region of AstaPo1. **A**. Structural formulae of several carotenoids which can be bound by AstaPo1. AXT – astaxanthin, CAN – canthaxanthin, ZEA – zeaxanthin, βCar – β-carotene. **B**. The domain structure of AstaPo1 showing the location of the predicted FAS1-like domain and the flanking N- and C-terminal regions. Note that the AstaPo1 preprotein contains an N-terminal signal peptide (residues 1-20), which was omitted in our construct as this peptide hampers the expression and purification of the soluble protein. Instead, the N terminus of the protein carried artificial residues ^17^GPHM^20^ left after the His_6_-tag removal. **C, D.** Analytical SEC profiles with diode-array detection, revealing the effect of AstaPo1 truncations on the size of the protein monomer (C) and its absorbance spectrum (D). The apparent *M*_w_ values determined from column calibration are indicated in kDa. **E**. Intrinsic Trp fluorescence spectra of AstaPo1 showing the effect of ZEA binding to the wild-type and truncated AstaPo1. Excitation wavelength was 297 nm. **F**. DSC thermograms showing the effect of carotenoid binding and truncation of the N- and C-terminal regions on the thermal stability of AstaPo1. *T*_m_ values are indicated as determined from the maxima of the peaks. Heating rate was 1 °C/min. The insert shows the color of the AstaPo1 samples with ZEA or CAN used.

To fill these gaps, here we study the AstaPo1 structure in complex with carotenoid and structure-function relationships, which provide insights into the carotenoid capture mechanism and inform the sequence-based prediction of the neofunctionalized carotenoid-binding ability in AstaP homologs.

## 2. Results

### 2.1. Localization of the ligand binding site within AstaPo1

We first wanted to localize the carotenoid-binding region within the mature AstaPo1 (residues 21-223, no signal peptide). Considering the presumable domain structure of this protein (**Fig. 1B**), we prepared AstaPo1 mutants lacking either the N-terminal region, or the C-terminal region, or both, and produced them in ZEA-synthesizing *E. coli* cells, which are known to yield stable ZEA-bound non-truncated AstaPo1 (17). Analytical SEC with continuous absorbance spectrum detection confirmed that truncations lead to a stepwise reduction of the protein monomer size (**Fig. 1C**), but do not affect the absorbance spectrum in the visible region. It retained the intact vibronic structure of the ZEA-bound AstaPo1 (**Fig. 1D**) even in the case of the shortest variant corresponding to the central domain, residues 41-190. A high Vis/UV absorbance ratio indicated unaltered and highly efficient carotenoid binding. Both 21-190 and 41-190 variants of AstaPo1 had the most pronounced Vis/UV ratio (~4, equal for both variants), which apparently reflected the removal of C-terminal Trp residues, W197 and W201 (**Fig. 1B**).

**Fig. 1E** shows that ZEA binding affected the intrinsic Trp fluorescence of AstaPo1. Having three Trp residues (W79, W197, W201), the AstaPo1 apoform exhibited a high-intensity Trp fluorescence, which was significantly quenched in the ZEA-bound form. Interestingly, the fluorescence maximum remained unchanged at 347 nm, which corresponds to partially exposed Trp residues (**Fig. 1E**) (18). Steady-state Trp fluorescence of the shortest ZEA-bound AstaPo1 variant, residues 41-190, which corresponds to the tentative FAS1-like domain containing a single Trp79 residue, was completely quenched (**Fig. 1E**), which confirms the direct carotenoid interaction within this domain. Given that Trp residues of carotenoproteins often form direct contacts with carotenoids (19–21), one may expect a similar involvement of Trp79.

Using DSC we found that the AstaPo1 apoform had a rather high denaturation temperature (*T*_m_ ~ 60 °C), whereas carotenoid binding substantially improved it further (to *T*_m_ of 75-80 °C) (**Fig. 1F**). Most importantly, truncation of the N- and C-terminal domains did not lower the high *T*_m_ for the AstaPo1 holoform (**Fig. 1F**).

Collectively, our data implied that the carotenoid-binding region of AstaPo1 is contained within its predicted FAS1 domain. Of note, while the apoform of the shortest AstaPo1 variant (41-190) corresponding to the FAS1 domain was entirely found in inclusion bodies, its holoform could be obtained upon expression in ZEA-producing *E. coli* cells. This suggests that the N- and C-terminal regions of AstaPo1 and carotenoid binding dramatically improve protein solubility. Successful formation of the AstaPo1^41-190^ holoform in *E. coli* cells indicates that such miniaturized AstaPo1 variant retains the ability to extract carotenoids from membranes in the absence of the N- and C-terminal regions, which are therefore not essential for this function. Given the tiny size of this water-soluble protein (16 kDa), it may be promising for various biomedical applications.

Nevertheless, the contributions from the tail regions to carotenoid embedment by AstaPo1 remained to be clarified by structural investigation in the context of the non-truncated protein.

### 2.2. Solution structure of AstaPo1 in complex with carotenoid

To investigate the structure of the AstaPo1 complex with its native carotenoid, AXT, we took advantage of solution NMR spectroscopy. We synthesized the ^13^C/^15^N-labeled AstaPo1 apoprotein and complexed it with the unlabeled chemically pure AXT. After extensive screening, we found the conditions allowing us to obtain 200-300 μM AstaPo1(AXT) samples and acquire the high-quality NMR spectra (**Supplementary Fig. S1**).

NMR chemical shifts of AXT inside the protein were elucidated using the isotope-filtered NOESY experiments (22). Such analysis revealed the presence of four sets of NMR signals, compared to the spectra of AXT acquired in organic solvents (**Supplementary Fig. S2**). One additional set of signals may arise due to the protein-induced asymmetry of the local environment around the two initially symmetrical parts of the AXT molecule (**Fig. 2A**). The other signals are most likely the consequence of conformational heterogeneity, which is in agreement with peak splittings observed in the protein NMR spectra (**Supplementary Fig. S3**). We analyzed several options that could explain this heterogeneity, including the alternative orientation of AXT within the cavity, cis-trans isomerism of double bonds (23), or the R-S isomerism of AXT, which is observed with respect to the position 3 (3’) of the β-ionone rings (24), and revealed that the latter phenomenon does actually take place (**Supplementary Text 1, Supplementary Figs S4–S6**). Therefore, we studied the spatial structure of AstaPo1(AXT) complex only for one of the states, (3S,3’S)-AXT, since the basic structural data for this isomer may be extracted from the PDB (25, 26).

**Fig. 2.**
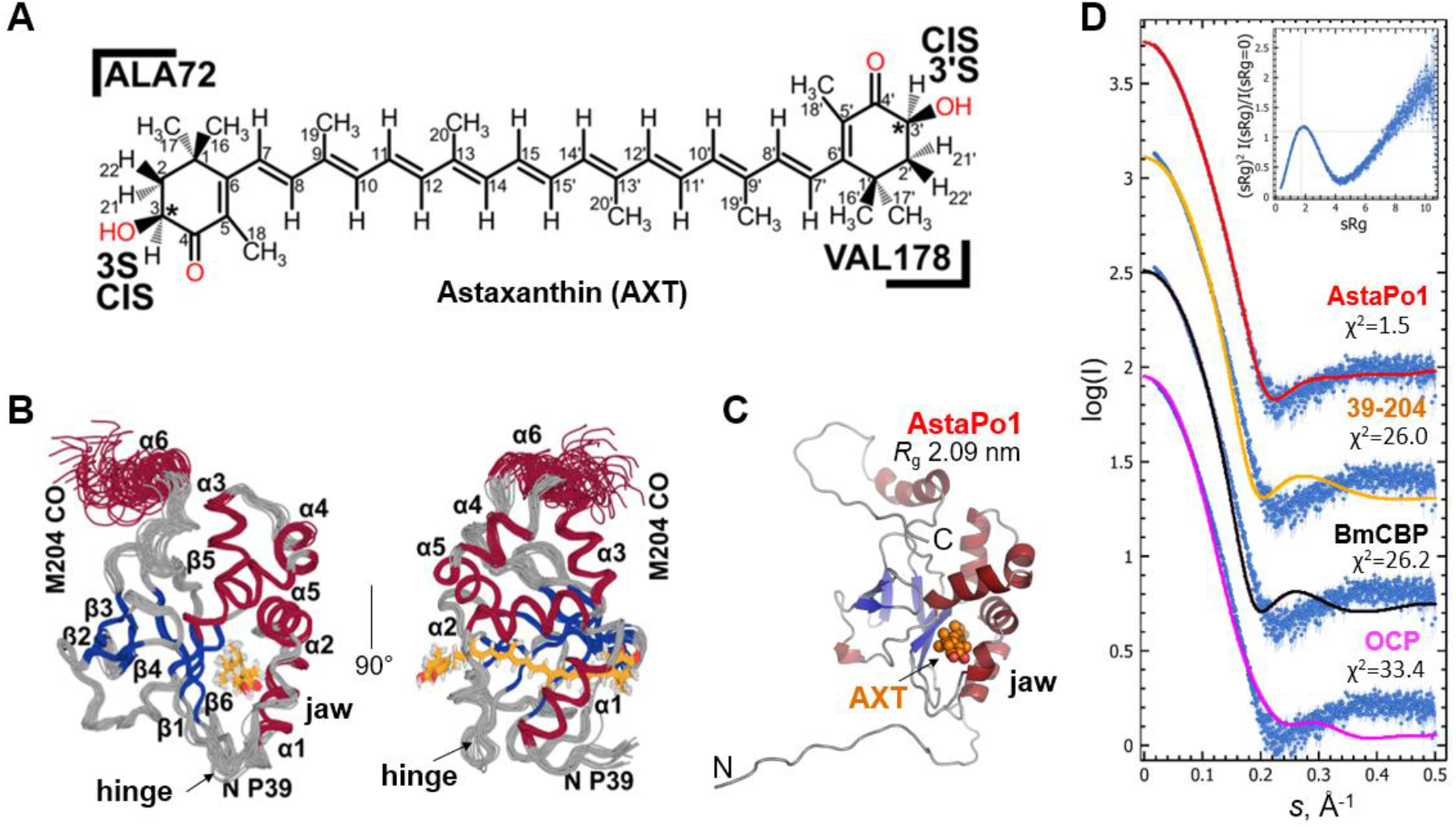
Spatial structure of the AstaPo1(AXT) complex. **A.** Chemical structure of the AXT stereoisomer used for NMR structure determination, with the indication of atom numbering. Asterisks indicate the chiral centers. Oxygenated groups are in red. **B.** Twenty NMR structures of AstaPo1(AXT) complex with the lowest restraint violations, shown superimposed over the backbone atoms of secondary structure elements (except for the C-terminal helix a6, residues 191-204) in two orthogonal views. The unstructured tails are hidden for clarity. Elements of the secondary structure are labeled. The AXT molecule is shown by orange sticks. **C.** A single representative full-atom NMR structure of AstaPo1(AXT) provides an excellent fit to the SAXS data collected for the AstaPo1(ZEA) complex (*R*_g_ of the model is 2.09 nm, X^2^ = 1.5; at 7 mg/mL). *R*_g_ of the twenty NMR full-atom structural models ranged 1.90-2.56 nm. Secondary structures are color-coded (red – a-helices, blue – β-strands, grey – unstructured regions). **D.** The fits to the SAXS data from the best-fitting NMR structure of the full-atom AstaPo1(AXT) complex, from the AstaPo1 FAS1 domain (residues 39-204; model *R*_g_ 1.85 nm), from the *Bombyx mori* Carotenoid-Binding Protein (model *R*_g_ 1.85 nm) (21), and from the *Synechocystis* Orange Carotenoid Protein (model *R*_g_ 2.05 nm) (27). The quality of the fits is indicated as X^2^ for each case. The insert shows the dimensionless Kratky plot for the AstaPo1(ZEA) complex. Dashed lines show the position of the maximum for the rigid sphere, which is given for reference.

Following the manual analysis of all the intermolecular NMR contacts, we found 150 intermolecular distance restraints and resolved the spatial structure of the AstaPo1(AXT) complex (**Fig. 2B, Table 1**). The central part of the protein adopts a typical fold of a FAS1 domain (3, 5) with a wedge-shaped β-sandwich, consisting of two three-stranded β-sheets and six a-helices. While five helices are typical for the FAS1 fold, the sixth, rather solvent-exposed C-terminal helix is peculiar in not being well packed into the globule with the other secondary structure elements, although it is anchored by the π-cation and ionic interactions between the W197 and E200 residues and charged side chains of a3 helix. Using the NMR relaxation parameters, we analyzed the dynamics of the AstaPo1 backbone and revealed that helix a6 tumbles as a whole with the protein, with some local mobility observed only at its C-terminal part (**Fig. S7–S8**). In contrast, we observed high-amplitude fast motions for the N- and C-terminal regions (residues 17-32 and 204-223) flanking the FAS1 domain. The structure of these parts of AstaPo1 is not well-defined in the NMR set, suggesting that the terminal residues are disordered, which is in line with the idea that they are chiefly required to maintain protein solubility.

**Table 1.** NMR input data and statistics for the AstaPo1(AXT) complex.

| *NMR distance and dihedral constraints* |  |
| --- | --- |
| Distance constraints |  |
| Total NOE for AstaPo1 | 1671 |
| Intra-residue | 447 |
| Inter-residue | 1224 |
| Sequential ($\vert i - j\vert = 1$) | 481 |
| Medium-range ($\vert i - j\vert \leq 4$) | 283 |
| Long-range ($\vert i - j\vert > 5$) | 460 |
| Intermolecular NOE | 150 |
| Hydrogen bonds | 52 |
| Total dihedral angle restraints |  |
| $\phi$ | 151 |
| $\psi$ | 155 |
| $\chi_1$ | 41 |
| Constraints per residue (excluding unstructured parts) | 15.1 |
| *Structure statistics (for set of 20 best structures)* |  |
| CYANA target function | 1.62±0.16 |
| Violations (mean±SD) |  |
| Distance mean[max], (Å) | 0.0056±0.0009 [0.23±0.07] |
| Dihedral angles mean [max], (°) | 0.382±0.042 [2.62±0.74] |
| RMSD of atom coords. ( stable secondary structure) (mean±SD) |  |
| Backbone (Å) | 0.57±0.04 |
| Heavy (Å) | 1.32±0.10 |
| RMSD of atom coords., AXT, (Å) | 0.31 ±0.08 |
| Ramachandran plot outliers |  |
| Residues in most favored regions | 94 % |
| Residues in additionally allowed regions | 6 % |
| Residues in generously allowed regions | 0 % |
| Residues in disallowed regions | 0% |
| *PDB ID* | **8C18** |

To justify the NMR structure, we exploited small-angle X-ray scattering (SAXS) of AstaPo1 complexes with either ZEA or CAN, as these particular holoproteins could be most efficiently produced in *E. coli* expressing the corresponding carotenoids. Despite the two distinct carotenoid types, differences in protein concentration, and variances in the residual amount of the apoform present in the samples (**Fig. 1F**), the SAXS data for AstaPo1(ZEA) and AstaPo1(CAN) yielded very similar structural parameters in solution (**Table 2**). Therefore, the type of bound carotenoid does not affect the protein conformation, and the SAXS data for the least noisy AstaPo1(ZEA) SAXS curve is representative also for AstaPo1(AXT). One of the NMR models provided an excellent fit (X^2^=1.5) to the SAXS data without any additional assumptions, whereas structural models of the individual FAS1 domain or of the other carotenoproteins of similar size (*R*_g_ ~2 nm) provided inadequate fits (**Fig. 2C-D**). Of note, the dimensionless Kratky plot for AstaPo1(ZEA) revealed the characteristic bell-shape and a gradual rise of the curve at large s\**R*_g_ values (**Fig. 2D**, insert), which reflected a nearly spherical folded protein with a number of peripheral flexible regions. Such description perfectly agrees with our NMR data.

**Table 2.** SAXS-derived structural parameters of the two AstaPo1 holoforms.

| Parameter | AstaPo1(ZEA) | AstaPo1(CAN) |
| --- | --- | --- |
| $M_w$ calculated from sequence | 21.6 kDa (apo) | 21.6 kDa (apo) |
| Number of residues | 207 | 207 |
| Protein concentration | 7.05 mg/ml | 2.57 mg/ml |
| $R_g$ (Guinier) | $2.09 \pm 0.01$ nm* | $2.13 \pm 0.01$ nm |
| $sR_g$ limits | 0.39-1.29 | 0.32-1.29 |
| $R_g$ (reverse) | $2.11 \pm 0.03$ nm | $2.18 \pm 0.04$ nm |
| $D_{max}$ | 7.8 nm | 8.1 nm |
| Porod volume | $38.1 \text{ nm}^3$ | $40.8 \text{ nm}^3$ |
| $M_w$ Porod | 23.8 kDa | 25.5 kDa |
| $M_w$ MoW | 14.3 kDa | 15.5 kDa |
| $M_w$ Vc | 21.1 kDa | 22.2 kDa |
| $M_w$ (size and shape) | 23.2 kDa | 25.3 kDa |
| Kratky plot | bell-shaped (folded) | bell-shaped (folded) |

\*All SAXS-derived parameters were calculated using the ATSAS 2.8 software package (45).

### 2.3. Carotenoid binding mode and protein-pigment interactions

NMR data revealed that AXT binds to AstaPo1 at the β-sheet composed of β1-β6-β5 strands and is covered by a jaw-like structure formed by two N-terminal helices a1, a2, and the a2-β1 hinge loop (**Fig. 2B and 3A**). In the observed binding mode, the hydrophobic carotenoid polyene is packed against the apolar side chains decorating the tunnel (at least 14 Ile/Val/Leu residues) (**Fig. 3A-E**), many of which are well conserved and belong to the FAS1-specific H1, H2 and YH motifs (**Fig. 3F**). At the same time, the β-ionone rings of the carotenoid protrude from the protein globule and are solvent-exposed, which explains the lack of fine structure in the absorption spectra of AstaPo1-bound AXT or CAN (**Fig. 3B,E**) (17). Surprisingly, we did not observe any specific protein-pigment interactions, such as hydrogen bonds or π-stacking found in other carotenoproteins with the known structure (19–21, 25). The only transient polar contact may take place between the side chain of Q56 and hydroxyl or carbonyl groups of AXT (**Fig. 3C,D**). Obviously, this contact cannot be formed in the case of β-carotene, one of the established AstaPo1 ligands (17). Hence, Q56 is perhaps not a strong carotenoid-binding determinant, although it could explain the apparently more efficient AstaPo1 binding with xanthophylls (17). While this residue is highly variable in the FAS1 domain-containing proteins (**Fig. 3F**), its chemical functionality is clearly preferred in AstaP orthologs with the proposed carotenoid-binding capacity (13–16) (see also below).

**Fig. 3.**
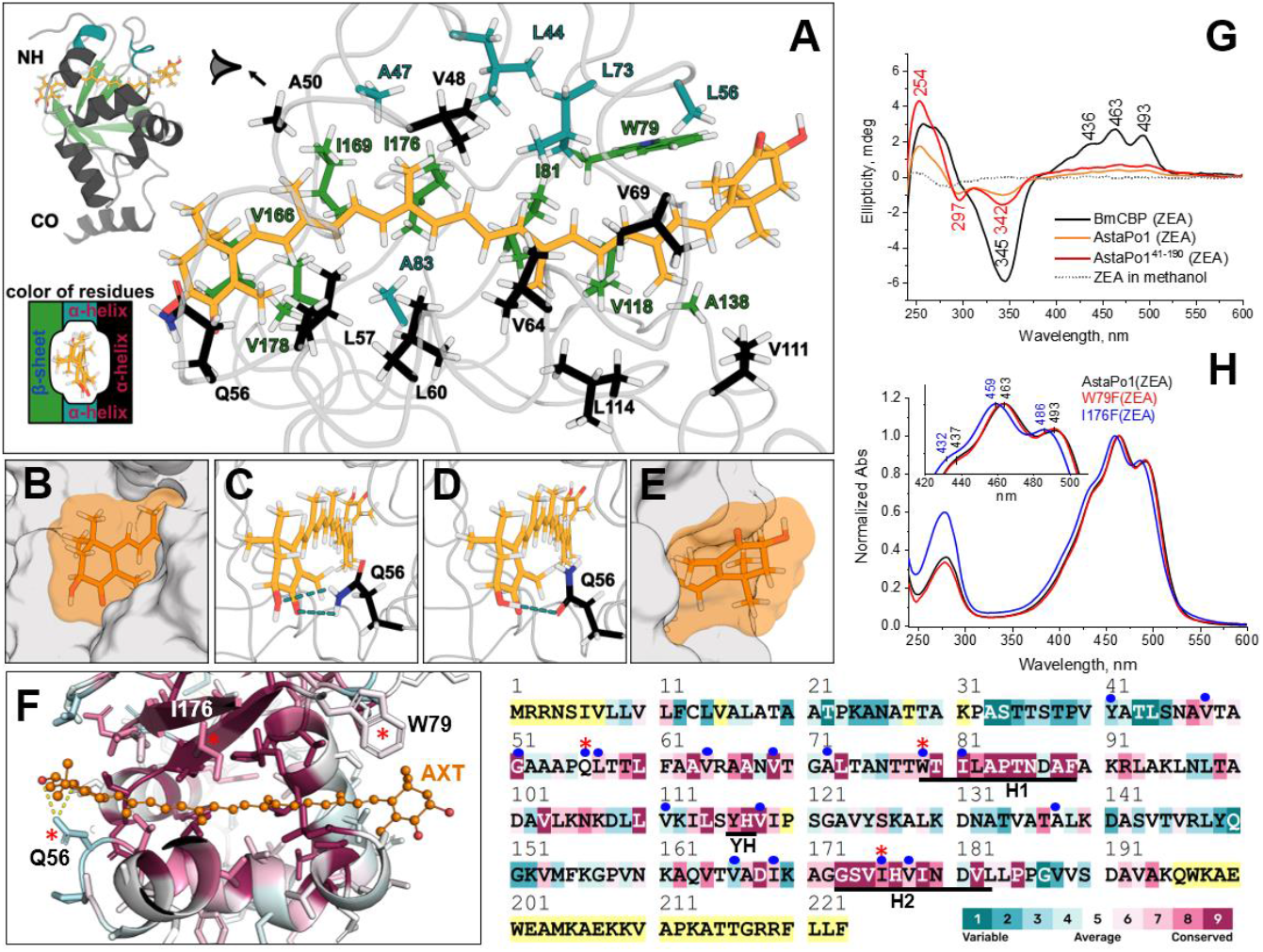
AXT binding mode and AstaPo1-pigment interactions. **A-E.** Different views of the carotenoid binding tunnel of AstaPo1. The coloring scheme in panel A is indicated. **B, E.** Polar headgroups of AXT exiting the protein molecule from both sides of the carotenoid-binding tunnel. The Connolly surface with solvent radius 1.4 A of AstaPo1 and the carotenoid is shown in grey and orange, respectively. **C, D**. Polar contacts formed between the AXT polar groups and the side chain of the non-conserved Q56 residue of AstaPo1. **F.** Amino acid conservation mapped onto the tertiary (left) and primary structure of AstaPo1 (right), according to Consurf (29) analysis of 150 FAS1 domain-containing homologs using the scale shown in the bottom right corner (yellow color indicates positions with insufficient data). The full AstaPo1 sequence is presented for the numbering consistency (residues 1-223). Red asterisks mark the peculiar positions discussed in the text. Blue circles highlight 17 residues whose solvent-accessible surface area changes by at least 10 Å^2^ upon AXT binding. FAS1-specific conserved motifs H1, H2, and YH are indicated by black bars. **G.** Vis-UV CD spectra of AstaPo1(ZEA) and its truncated variant AstaPo1^41-190^(ZEA) as compared with the spectra of BmCBP(ZEA) and free ZEA in methanol. The main extrema are indicated in nm. **H.** Absorbance spectra of AstaPo1, its W79F and I176F variants purified from *E. coli* cells synthesizing ZEA.

We did not find any special contribution from the disordered tail regions into the interface with AXT, which supports the conclusions made above for the truncated AstaPo1. Since the main protein-pigment interactions revealed by the AstaPo1(AXT) structure are the hydrophobic contacts holding the polyene chain of the carotenoid in the conserved hydrophobic tunnel of the AstaPo1 structure, we identify no features that would determine the ligand specificity. Thus, the protein should be able to bind any sufficiently long rod-like hydrophobic molecules with the optional presence of polar headgroups.

According to the low-amplitude Vis/UV CD spectra, the carotenoid molecule in AstaPo1 is rather symmetric and straight, in contrast to the curved conformation previously reported for BmCBP (21) or OCP (28) carotenoproteins (**Fig. 3G**). This supports the notion that the AstaPo1-bound AXT does not experience any significant torsions, which is often a consequence of direct chemical contacts with the carotenoid rings, such as in OCP (19) or in β-crustacyanin (25). In nice agreement with the Trp fluorescence quenching data (**Fig. 1E**), our NMR structure reveals that one of the AXT rings neighbors the side chain of the single Trp79 in the carotenoid-binding domain of AstaPo1, although its conformation and distance (>5 A) from the ring are incompatible with H-bonds or π-stacking interactions that would stabilize the bound carotenoid (**Fig. 3A**). Interestingly, Trp in this position is not fully conserved in the FAS1 protein superfamily, and is often replaced by a Phe (**Fig. 3F**). Crucially, the amino acid substitution W79F in AstaPo1 produced in ZEA-synthesizing *E. coli* cells, does not abolish its ZEA-binding capacity nor change the absorbance spectrum (**Fig. 3H**), which supports the idea that the indolyl group in this position is dispensable for carotenoid binding. To test the possible importance of the tunnel residues, we also obtained the I176F mutant with a Phe residue introduced in the conserved H2 motif in the middle of the tunnel (**Fig. 3F**) and produced it in ZEA-synthesizing *E. coli* cells. While it surprisingly retained the carotenoid-binding ability, its efficiency was compromised (only 60% of the holoform), as judged from a substantially lowered Vis/UV absorbance ratio of 1.7 instead of 2.8 for the wild-type (**Fig. 3H**). Curiously, the Vis absorbance of ZEA bound to the I176F mutant was appreciably blue-shifted by ~4-7 nm, with an altered fine structure, compared with the wild-type AstaPo1 (**Fig. 3H**). Such spectral change reflects the renowned sensitivity of the carotenoid absorbance to even subtle changes in microenvironment (21) and the effect of the bulky Phe side chain in close vicinity of the carotenoid polyene.

### 2.4. Insights into the carotenoid capture mechanism

The apparent flexibility of the a2-β1 hinge loop (**Fig. 2B, S7**) implies that in the absence of the ligand, the jaw formed by a1-a2 helices likely samples some conformational space around its rather closed position in the AstaPo1(AXT) complex, which would facilitate ligand uptake. The a1-a2 jaw motions are indirectly supported by the NMR data obtained for the AstaPo1 apoprotein. The poor quality of NMR spectra and limited stability of the apoprotein did not allow direct structure determination. However, we managed to obtain the partial (63%) assignment of the NMR chemical shifts (**Fig. S9**), which covered the N- and C-terminal tails, helices a3, a4, a5 (partially) and a6, strands β2-β6 (**Fig. 4A**). According to the secondary chemical shifts, for the covered regions we observe no difference between the apo and holo states (**Fig. S10**), implying that the structure of β-sheet and helices distant from the ligand-binding site is preserved. On the other hand, the NMR signals of the a1-a2 jaw and adjacent loops are either not seen or substantially broadened, which indicates that this AstaPo1 part becomes mobile in the μs-ms timescale and indeed may detach from the hydrophobic core of the protein.

**Fig. 4.**
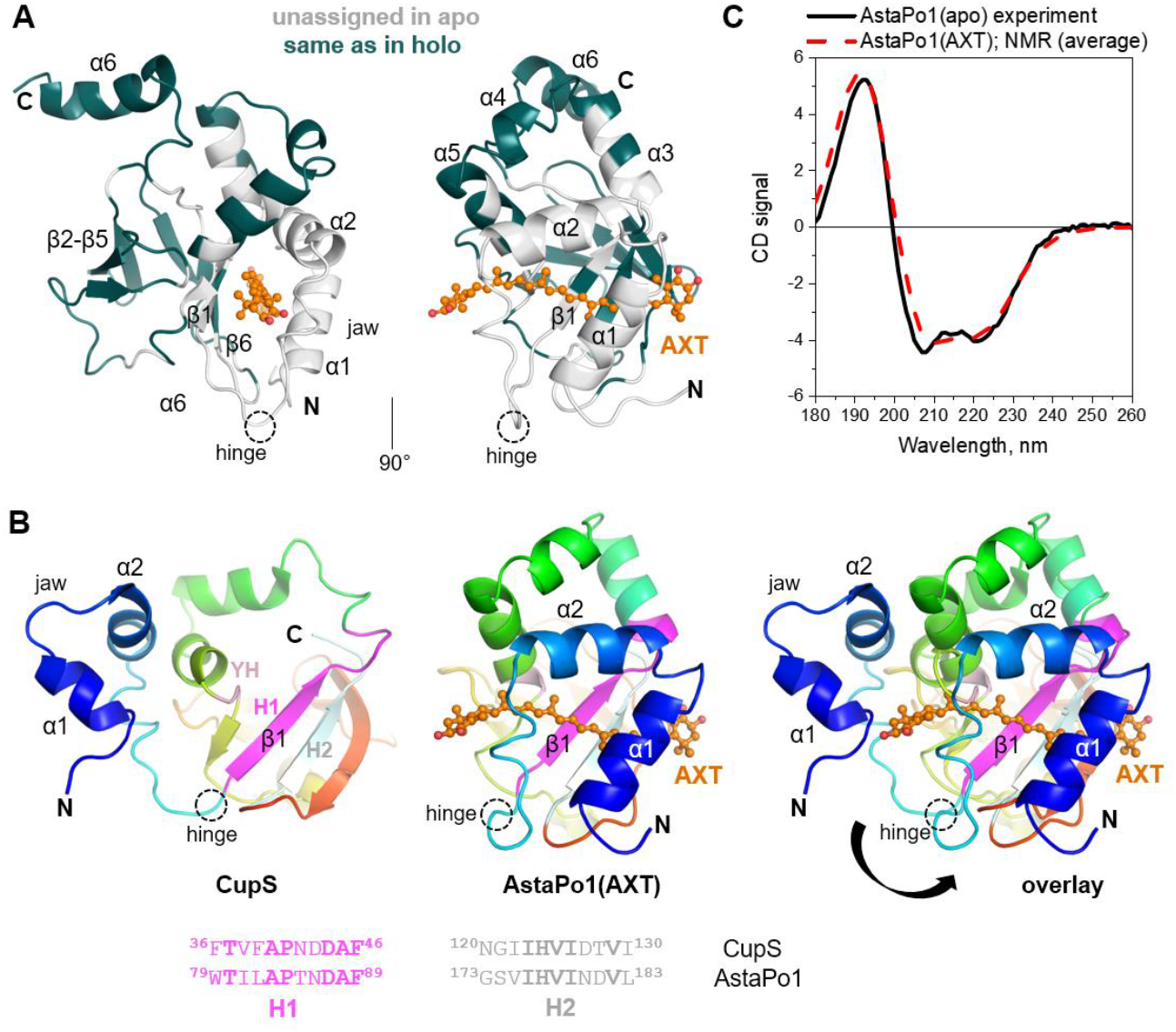
Insights into the carotenoid capture mechanism by AstaPo1. **A.** Structure of the AstaPo1/AXT complex is colored according to the NMR data obtained for the apoform. Regions with known NMR chemical shifts which reveal the identical structure with the AstaPo1 holoform are colored green (**Fig. S10**), and the other parts of the protein, for which no assignment could be obtained, are colored light grey. **B.** A hypothetical carotenoid capture mechanism. Three panels represent from left to right: the CupS FAS1-containing protein (PDB ID: 2MXA (7)) in a tentative “open-like” conformation, AstaPo1 in complex with AXT (our NMR structure) and their overlay. The structures are colored by a gradient from blue (N) to red (C), except for the conserved motifs used for structural alignment - H1 (magenta), H2 (pale cyan) and YH (pale pink). Pairwise sequence alignment of the H1 and H2 fragments is shown below (identical residues are in bold font). AXT is shown as an orange ball-and-stick model, the main secondary structure elements and the hinge loop are labeled. The proposed conformational transition from an open to the carotenoid-bound state is depicted by the arrow. **C.** Far UV CD spectrum of the AstaPo1 apoform (black line) shown overlaid with the average CD spectrum calculated from 20 NMR models of the holoform of the same construct (blue dashed line).

A survey of the published structures of the FAS1 domains helped to find a peculiar structure of the single-domain FAS1 protein CupS from *T. elongatus* (NDH-1 complex sensory subunit, Uniprot Q8DMA1: 30.8% sequence identity with AstaPo1 FAS1) (7) (**Fig. 4B**). When superimposed onto our AstaPo1(AXT) structure using the highly conserved H1/H2 motifs and YH dyads, the CupS structure is remarkably dissimilar by the position of the a1-a2 jaw, suggesting an imaginary rotation over the a2-β1 hinge loop which would resemble a conformational transition from an open-like to the closed, AXT-bound conformation. Of note, such transition would not involve significant secondary structure rearrangements. We confirmed this using far UV-CD spectroscopy of the apo- and carotenoid-bound forms of AstaPo1 (**Fig. 4C**). Due to the unavoidable contribution of carotenoid to the experimental CD spectrum of the AstaPo1 holoform, the latter was calculated from NMR models of AstaPo1(AXT) using PDBMD2CD (30). The similarity of the apo/holo CD spectra supports the idea of jaw movement as a whole. While remaining speculative, such rearrangement provides a useful insight into the carotenoid capture by AstaPo1.

Thus, we conclude that, while the apoform is not different from the AXT-bound state of AstaPo1 in terms of the overall secondary structure, the flexibility and loop conformations of the a1-a2 jaw and its surroundings change dramatically in response to the ligand uptake.

### 2.5. Carotenoid binding as a privilege function of a subset of FAS1 domain containing proteins

FAS1 domains have an evolutionarily ancient and wide-spread fold, however, the ligand binding capacity of such proteins has not been reported, until recently. Therefore, the possible structure and sequence determinants of such a new FAS1 function warranted analysis. Based on the NMR structure, we were able to identify AstaPo1 residues directly involved in carotenoid binding by calculating the solvent-accessible surface area (SASA) changes with and without carotenoid (using 10 A^2^ as cutoff) and map them onto the primary and tertiary structure (**Supplementary Fig. S11**). By comparing how these residues are conserved in FAS1 domain-containing proteins in general (**Fig. 3F**) and in tentative AstaP orthologs in *Scenedesmaceae* with the reported or expected carotenoid-binding capacity (13–16) (**Fig. 5A**), we revealed that many positions are occupied by identical or similar residues in both groups, namely, 48, 51, 57, 64, 69, 79, 81, 118, 169, 176, 178 (AstaPo1 residue numbering). Most of these residues decorate the carotenoid-binding tunnel of AstaPo1, which is preserved in many FAS1 members regardless of their prospective ligand-binding ability. Positions 41, 56, 72, and 166 are variable in both groups, which means that their chemical functionality is likely dispensable for carotenoid binding. Given their location on the periphery of the hydrophobic tunnel, these residues even in genuine carotenoid-binding AstaP orthologs likely play a secondary role in ligand binding, such as in the case of Q56, whose side chain forms transient H-bonds with AXT rings (**Fig. 3**), but obviously does not contribute to the binding of non-oxygenated β-carotene (17). While among the ten tentative carotenoid-binding FAS1 domains the glutamine is conserved at least in six, and in one more case is replaced by a synonymous Asn (**Fig. 5A**), its replacement by another polar residue cannot disqualify an AstaP homolog as a carotenoid binder a priori. Most intriguing, some carotenoid-contacting residues in AstaPo1 are identical or similar in AstaP orthologs but are not conserved throughout the FAS1 superfamily (namely, positions 111 and 138 at one of the tunnel exits). This favors the idea that the carotenoid-binding function is a feature of only a subset of such proteins.

**Fig. 5.**
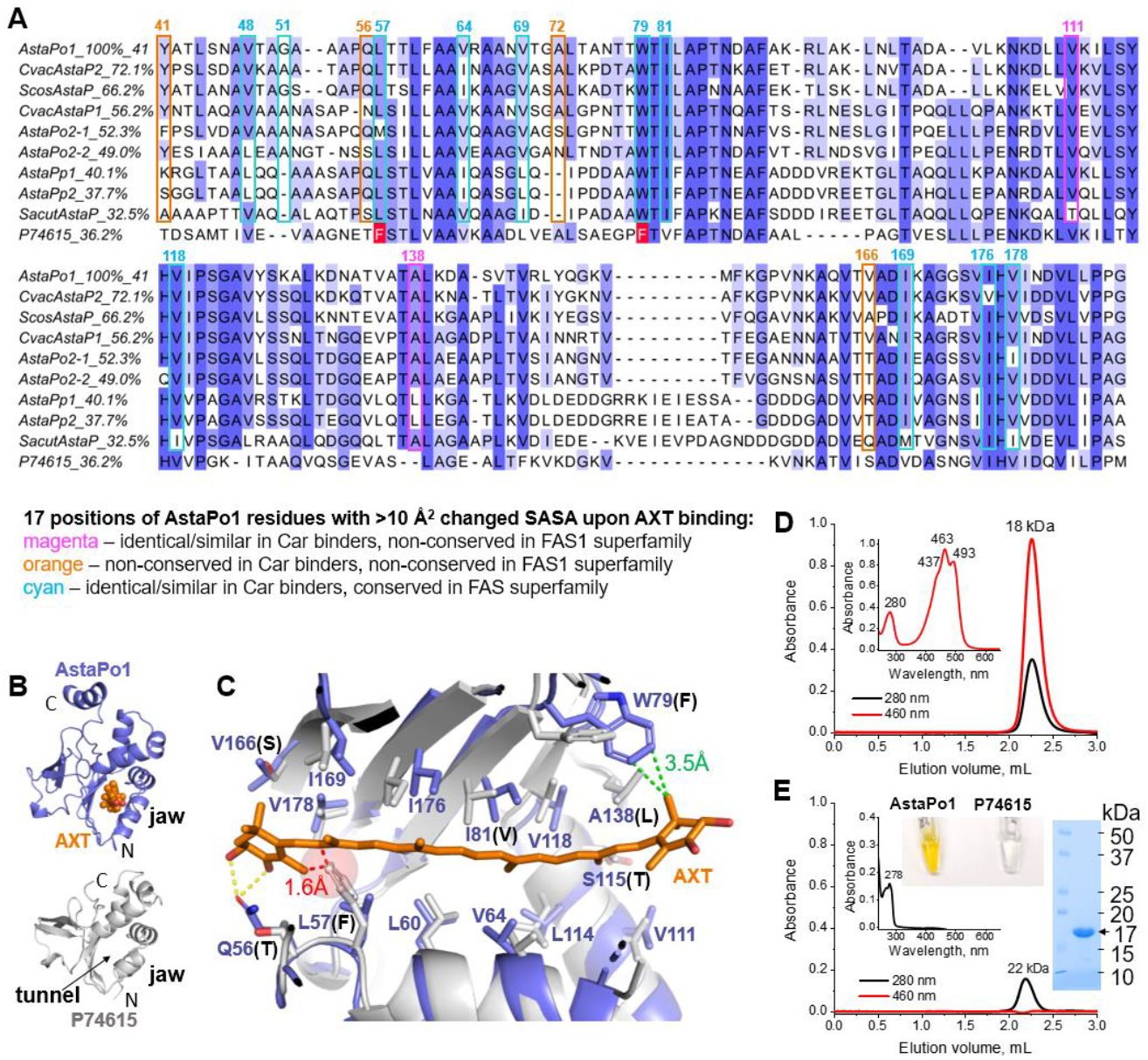
Carotenoid binding as neofunctionalization of AstaP orthologs. **A.** MSA of the FAS1 domains of the tentative carotenoid (Car) binding AstaP orthologs from *Coelastrella astaxanthina* (AstaPo1; BAN66287.1), *Coelastrella vacuolata* (CvacAstaP1, QYF06643.1; CvacAstaP2; QYF06644.1), *Scenedesmus costatus* (ScosAstaP, QYF06645.1), *Scenedesmus sp*. Oki-4N (AstaPo2-FAS1 domain 1 and AstaPo2-FAS1 domain 2, BBN91622.1; AstaPpink1, BBN91623.1; AstaPpink2, BBN91624.1), *Scenedesmus acutus* (SacutAstaP; ACB06751.1) and *Synechocystis* sp. PCC 6803 (SynAstaP, Uniprot P74615), shown with coloring by similarity as shades of blue. NCBI accession numbers are indicated for each sequence (except SynAstaP) in parentheses. Sequence identity of each ortholog relative to AstaPo1 is indicated in %. Seventeen positions of the AstaPo1 sequence whose solvent accessible surface area (SASA) changed > 10 A^2^ upon AXT binding (NMR data) are mapped on the MSA to reveal positions satisfying the criteria indicated by magenta, orange and cyan. Note that positions of Leu57 and Trp79 of AstaPo1 are occupied by Phe residues in SynAstaP (P74615) (marked by red). **B**. Comparison of the NMR structure of AstaPo1 complexed with AXT and the Alphafold2 model of *Synechocystis* ortholog P74615, showing the similar FAS1-like fold featuring the jaw and the tunnel. **C.** Structure alignment of AstaPo1(AXT) complex (blue) and the Alphafold2 model of P74615 indicating the nearly identical tunnel lining and different tunnel exits. Note that the side chain of the non-conserved Phe58 of P74615 clashes with the carotenoid polyene, that the non-conserved Gln56 of AstaPo1 contacting the polar groups of AXT is replaced by a shorter Thr56 residue in P74615, and that the nearly absolutely conserved Trp79 of AstaPo1 is replaced by Phe80 in P74615 (the corresponding distances are shown). **D, E.** The zeaxanthin-binding capacity of AstaPo1 (50 μM) and P74615 (500 μM) analyzed by SEC of proteins purified from *E. coli* cells synthesizing zeaxanthin. Note that P74615 has an extremely low extinction coefficient at 280 nm due to the total absence of Trp and presence of only one Tyr residue, hence was loaded in the 10-fold higher concentration. The apparent *M*_w_ of the peaks are shown. The absorbance spectra recorded during SEC runs and corresponding to the peak maxima are presented in the inserts. Absorbance maxima are indicated. The inserts in panel D also show the electrophoretic purity of the P74615 preparation used for the analysis and the appearance of the AstaPo1 and P74615 samples purified from ZEA-synthesizing *E. coli* cells.

A relatively distant AstaPo1 homolog from the cyanobacterium *Synechocystis* sp. PCC 6803 (Uniprot ID: P74615, sequence identity with AstaPo1 - 35.75%) with the anticipated carotenoid-binding capacity is especially remarkable (**Fig. 5A**) (13). Among the 17 carotenoid-contacting positions in AstaPo1, the P74615 sequence markedly differed by the presence of the TF cluster instead of the ^56^QL^57^ residues of AstaPo1, and also by the presence of the Phe residue instead of Trp79. The neutral effect of the W79F replacement on the carotenoid-binding ability of AstaPo1 (**Fig. 3H**) likely qualifies the corresponding sequence difference as non-crucial. The secondary role of Q56 has been discussed above. Meanwhile, the Leu→Phe substitution at the generally conserved position 57 (**Fig. 3F, 5A**) prompted us to perform the structural comparison. The Uniprot P74615 entry annotates a protein with the predicted C-terminal FAS1 domain (residues 45-176), and its Alphafold2 model reveals a striking similarity with our AstaPo1 structure (Cα RMSD of 1.8 Å) (**Fig 5B**). The hydrophobic tunnels of the two proteins are nearly identical, yet the Phe58 residue of the TF dyad in the P74615 homolog is appreciably bigger than an equivalent Leu57 residue in AstaPo1 (**Fig. 5C**). Interestingly, a Phe in this position is present in as many as 88 out of 150 FAS1 homologs used for our Consurf analysis (**Fig. 3F**), while a Leu residue is present in only 23 of 150. Hypothesizing that such a bulky residue could interfere with the carotenoid binding, we produced the P74615 protein in ZEA-synthesizing *E. coli* cells to compare its carotenoid-binding capacity with that of AstaPo1. The P74615 protein could be well expressed and formed a soluble monomer (apparent *M*_W_ ~22 kDa). However, under the conditions when AstaPo1 efficiently bound ZEA acquiring the characteristic absorbance spectrum (**Fig. 5D**), the P74615 protein could be obtained only as an apoprotein, which lacked any absorbance in the visible spectral region (**Fig. 5E**). This example supports the idea that it was variations of the peripheral residues at the exits of the hydrophobic tunnel that may have determined the newly acquired ability of some FAS1 members to bind ligands.

## 3. Discussion

In this work, we determined the first NMR structure of a carotenoid-protein complex, which, together with the biochemical data, revealed the peculiar mechanism of carotenoid embedment and explained the lack of AstaP specificity to carotenoids (17). The determined structure is unique for carotenoid-binding proteins and supports the idea that, being found in many different organisms, carotenoid-binding proteins emanate from the convergent evolution - their spatial organization, the mechanism of carotenoid uptake, and the ligand specificity are completely different.

For example, perhaps the earliest structurally described, β-crustacyanin is a 42-kDa heterodimer specifically binding two AXT molecules and controlling the coloration of lobster shells (25). The Orange Carotenoid Protein (OCP) is a two-domain, 35 kDa photoswitching protein that binds a single ketocarotenoid molecule and regulates photoprotection in cyanobacteria (19). The independently existing homologs of the N- and C-terminal domains of OCP, widely distributed among cyanobacteria, are carotenoid-binding proteins that participate in interprotein carotenoid transfer processes (31–33). The structure of the 18 kDa C-terminal domain homolog (CTDH) has been determined by X-ray crystallography and NMR in the apoform only (20, 34), but it is known to homodimerize upon carotenoid binding (28, 33, 35). Being a rather specific ketocarotenoid binder, CTDH proved to be a robust carotenoid delivery module which dissociates to monomers upon donating the carotenoid to other proteins or to biological membrane models (36). In contrast, the homologs of the N-terminal domain of OCP, the so-called Helical Carotenoid Proteins, form rather small all-α helical ~20 kDa monomers, which are less specific and can bind at least deoxymyxoxanthophyll, echinenone, canthaxanthin and β-carotene (31). Despite a broader ligand repertoire, HCP is a poorly efficient carotenoid extractor - it has always been purified along with the large excess of the apoform and required a CTDH homolog for the carotenoid delivery and maturation into the holoform (32, 37). The recently reported crystal structure of the 27 kDa Carotenoid-Binding Protein from silkworm (BmCBP) (21), which determines coloration of the silkworm cocoons (38), revealed the STARD3-like fold accommodating only part of the bound carotenoid molecule in the lipid-binding cavity (39, 40). This protein can bind a wide variety of carotenoids and can transfer them to other proteins and liposomes, representing an attractive carotenoi-ddelivery module for various biotechnological and biomedical applications (41).

The AstaPo1 holoform structure determined here (**Fig. 2**) is dissimilar to any of these carotenoproteins. We show that its central domain has a FAS1-like fold, but the wide superfamily of FAS1-containing proteins has no common ligand-binding functions reported, not to mention that AstaP was discovered as the first FAS1 protein binding carotenoids (13). Looking at the FAS1 fold it was nearly impossible to guess the exact location of the carotenoid-binding site *a priori*, especially since the homolog studied, AstaPo1, has rather long regions flanking the FAS1 domain, with the undefined structure and unclear contribution to carotenoid binding. We first localized the carotenoid-binding site to the FAS1 domain and showed that such miniaturized AstaPo1 (as small as 16 kDa) successfully matured into the holoform upon expression in carotenoid-producing *E. coli* strains, and is one of the smallest proteinaceous carotenoid delivery modules currently known. Since the tailless apoform was fully insoluble, the tails, variable in amino acid composition and length in AstaP orthologs (14, 15), are likely required for maintaining protein solubility, which is especially important given that this protein is up-regulated under stress conditions and functions at high concentrations. We anticipate that the AstaPo1^41-190^ variant can be shortened even further without the loss of the carotenoid-carrying capacity.

Our NMR and mutagenesis data indicate that carotenoid binding involves a tunnel decorated by highly conserved hydrophobic amino acids, but its length is insufficient for accommodating the entire 30-A carotenoid molecule. Thereby the carotenoid rings protrude from the globule and experience no apparent specificity restrictions, whereas the carotenoid polyene is stably fixed in the hydrophobic tunnel (**Fig. 3**). Perhaps, the only specific interactions with the carotenoid rings observed in the NMR structure are H-bonds involving the Gln56 side chain on the periphery of the tunnel. Although these interactions obviously appear favoring for binding xanthophylls, they should have no role in the binding of β-carotene, and hence Gln56 is unlikely a strict carotenoid-binding determinant. Yet, the majority of AstaP orthologs with the reported or proposed carotenoid-binding function have this Gln (or its replacement by an Asn) (**Fig. 5**). Intriguingly, this position is not conserved at all in the context of the entire FAS1 superfamily, which is also the case for some other positions occupied by the carotenoid-contacting residues in our structure (**Fig. 5**). The analysis of covariation of such residues revealed that positions on the periphery of the tunnel can favor or instead disfavor the carotenoid-binding capacity, which informs the sequence-based prediction of such function in AstaP homologs. To illustrate the practical usefulness of such prediction, we selected a cyanobacterial AstaP homolog with a similar domain structure but a remarkable, unfavorable replacement of the Leu57 residue of AstaPo1 by a bulky Phe, and experimentally demonstrated that it indeed cannot bind carotenoids, in striking contrast to AstaPo1 (**Fig. 5**). It is especially intriguing that the Phe residue is found in as many as 88 out of 150 FAS1 sequences used in our evolutionary analysis (**Fig. 3**).

Since cell adhesion principles involving FAS1 domains are thought to have evolved at the earliest known stages of evolution (2), the carotenoid-binding function is likely an example of neofunctionalization of a subset of AstaP-like proteins within the green algae. Finding and experimentally validating carotenoid-binding orthologs of AstaPo1 warrant further interesting investigations.

## 4. Materials and methods

### 4.1. Materials

All-trans-astaxanthin (CAS Numbers: 472-61-7) was purchased from Sigma-Aldrich (USA). Absorbance spectra of carotenoids in organic solvents were registered on a Nanophotometer NP80 (Implen, Germany) using the following molar extinction coefficients: 125,000 M^-1^ cm^-1^ for AXT at 482 nm in DMSO (42), 145,000 M^-1^ cm^-1^ for ZEA at 450 nm in methanol (43). Chemicals were of the highest quality and purity available.

### 4.2. Plasmid construction and mutagenesis

AstaPo1 cDNA corresponding to residues 21-223 of the Uniprot S6BQ14 entry was codon-optimized for expression in *E. coli*, synthesized by Integrated DNA Technologies (Coralville, Iowa, USA) and cloned into the pET28-His-3C vector (kanamycin resistance) using the *NdeI* and *XhoI* restriction sites. SynAstaP cDNA corresponding to residues 27-180 of the Uniprot P74615 entry was codon-optimized for expression in *E. coli*, synthesized by Kloning Fasiliti (Moscow, Russia) and cloned as described above.

AstaPo1 truncated variants (AstaPo1^21-190^, AstaPo1^41-223^, AstaPo1^41-190^) or mutants W79F and I176F were obtained by the overhang PCR or by the megaprimer PCR method respectively using Q5 (NEB) polymerase, primers listed in **Supplementary Table S1**. The same vector and restriction sites as above were used. The N terminus of all proteins contained extra residues GPHM after 3C cleavage step. All resulting constructs were verified by DNA sequencing (Eurogen, Moscow, Russia).

### 4.3. Protein synthesis and sample preparation

The wild-type AstaPo1, its mutant constructs, and SynAstaP were transformed into BL21(DE3) *E. coli* cells for expression of the apoforms. For the production of carotenoid-bound holoforms plasmids described above were transformed into either BL21(DE3), carrying pACCAR25ΔcrtX plasmid (chloramphenicol resistance, ZEA biosynthesis), or BL21(DE3), containing pACCAR16ΔcrtX (chloramphenicol resistance, beta-carotene biosynthesis) and the pBAD plasmid (ampicillin resistance, CAN biosynthesis). Details on the pathway of carotenoid biosynthesis in our system are described in (17).

Expression cultures were grown in LB medium till OD_600_ = 0.6 at 37 °C and induced by 0.05-0.2 mM IPTG for 24 hours at 25 °C. In the case of wild-type holoforms, expression was carried at 30 °C and for CAN biosynthesis addition of 0.02% L-arabinose was also required. For expression of AstaPo1 mutant holoforms temperature was decreased to 25 °C.

All recombinant proteins were purified using the combination of subtractive immobilized metal-affinity and size-exclusion chromatography to electrophoretic homogeneity. Pure proteins were aliquoted and stored frozen at −80 °C. Protein concentrations were determined on a Nanophotometer NP80 (Implen) using the extinction coefficients given in **Supplementary Table S2**. For AstaP holoforms, protein extinction coefficients were used after subtraction of the carotenoid contribution into absorbance at 280 nm determined earlier, which are *A*_463_/6 for ZEA and *A*_479_/7.4 for CAN (17). The expected Vis/UV absorbance ratios for AstaPo1 complexes with ZEA are listed in **Supplementary Table S2**. The carotenoid content of holoforms was analyzed by acetone-hexane extraction of carotenoids followed by thin-layer chromatography (41).

For NMR, the His_6_-tagged AstaPo1 was synthesized in *E. coli* BL21(DE3) cells. Overnight grown *E. coli* were diluted till OD_600_=0.005 in 2 L of fresh M9 minimal salt medium supplemented with Traces of metals (1:10000 v:v (44)) and containing 50 mg/mL kanamycin. For isotopic labeling, ^15^NH_4_Cl and ^13^C D-glucose were used. Cultures were grown to OD_600_=0.6 at 28 °C and then the protein expression was induced by the addition of 0.1 mM IPTG for 24 h incubation at 16 °C.

Cells containing the over-expressed AstaPo1 were harvested at 7000 g, 4 °C for 5 min and resuspended in 120 ml of lysis buffer (0.1 M Tris pH 7.5; 1 M NaCl; 1 mM BME; 10 mM Imidazole; 10% glycerol; 1% Triton X-100 and 0.2 mM PMSF). Resuspended cells were disrupted by sonication for 30 cycles of 20 sec of sonication and 2.5 min of resting and the cell lysate was spun at 14,000 g, 4 °C for 1 h. The pellet was discarded and the supernatant was passed through a 0.22 μm filter (Millipore) and purified by affinity chromatography using a 5 mL Ni-NTA (Qiagen) column. Immobilized AstaPo1 was washed thrice, first with buffer containing 0.1 M Tris pH 7.5; 1 M NaCl; 1 mM BME; 0.5% Triton X-100 and 10 mM Imidazole, next with the buffer containing 0.1 M Tris pH 7.5; 1 M NaCl; 1 mM BME and 40 mM Imidazole and final washing step with buffer containing 0.1 M Tris pH 7.5; 1 M NaCl; 1 mM BME and 80 mM Imidazole. His-tagged AstaPo1 was eluted with a buffer containing 0.1 M Tris pH 7.5; 1 M NaCl; 1 mM BME and 500 mM imidazole. Eluted AstaPo1 was dialyzed overnight against 100x volume of 50 mM Tris buffer pH 7.5 containing 150 mM NaCl, in dialyzer D-Tube^™^ Dialyzer Mega, MWCO 3.5 kDa (Merck) at 4 °C.

Overnight incubation with 3C protease from human rhinovirus was used to remove the His6-tag of AstaPo1. Next, the sample was centrifuged at 25,000 g, 4 °C for 1 h and purified by affinity chromatography using a 5 ml Ni-HP column (Qiagen). Flow through with pure AstaPo1 was collected and dialyzed overnight against the buffer containing 50 mM Tris pH 7.6, 0.15 M NaCl in dialyzer D-Tube^™^ Dialyzer Mega, MWCO 3.5 kDa (Merck) at 4 °C. Pure AstaPo1 was used immediately or stored at −20 °C.

To assemble the AstaPo1/AXT complex, we mixed AXT in DMSO (1 mg/ml) with purified AstaPo1 at the 4.3-fold excess and incubated at moderate mixing, room temperature for 1h, light-protected. DMSO contents in the solution were 10-12%. All protein-free AXT was removed by centrifugation at 25,000 g, 4 °C for 1 h. The supernatant was dialyzed against the buffer containing 50 mM Tris pH 7.5 and 150 mM NaCl at 4 °C, overnight to remove the DMSO. The AstaPo1/AXT complex was concentrated by an Amicon concentrator with 3.5 kDa MWCO (Merck) to a concentration of 7 mg/mL (280 μM).

### 4.4. Analytical size-exclusion spectrochromatography

SEC with continuous diode-array detection in the 240-850 nm range (recorded with 1-nm steps (4 nm slit width) and a 5 Hz frequency) was applied to study the apparent Mw and absorbance spectra of the protein holoforms. Samples (50 μl) were loaded on a Superdex 200 Increase 5/150 column (GE Healthcare, Chicago, Illinois, USA) pre-equilibrated with a 20 mM Tris-HCl buffer, pH 7.5, containing 150 mM NaCl, 3 mM NaN_3_, 5 mM β-ME and operated using a Varian ProStar 335 system (Varian Inc., Melbourne, Australia). Diode-array data were converted into .csv files using a custom-built Python script and processed into contour plots using Origin 9.0 (Originlab, Northampton, MA, USA).

### 4.5. Circular dichroism

AstaPo1 (0.5 mg/ml, 23 μM, Vis/UV absorbance ratio 2.54) or its truncated AstaPo1^41-190^ mutant (0.42 mg/ml, 26.5 μM, Vis/UV absorbance ratio of 3.35) were dialyzed overnight against 20 mM Na-phosphate buffer pH 7.05 and centrifuged for 10 min at 4 °C and 14,200 g before measurements. Visible/UV CD spectra were recorded at 20 °C in the range of 190-650 nm at a rate of 0.4 nm/s with 1.0 nm steps in 0.1 cm quartz cuvette on a Chirascan circular dichroism spectrometer (Applied Photophysics) equipped with a temperature controller. The raw spectrum was buffer-subtracted before presentation. Free ZEA (12 μM) in methanol was measured for reference. ZEA concentration in methanol was determined spectrophotometrically. The Vis/UV CD spectrum of the BmCBP(ZEA) complex (0.5 mg/ml, 18.5 μM, Vis/UV absorbance ratio 1.42) was taken from previous work (21). For normalization, all CD spectra were scaled so that the corresponding absorbance spectra in the visible range were of the same amplitude.

To compare the secondary structures of the apo- and holoform of AstaPo1 we used the experimentally determined far UV CD spectrum of AstaPo1(apo) (17) and calculated the far UV CD spectrum of the AstaPo1(AXT) complex based on the NMR structures. To this end, we first calculated the CD spectra for each of the twenty NMR models in PDBMD2CD (30) and then scaled the average CD spectrum to that of the apoform for comparison.

### 4.6. Fluorescence spectroscopy

Steady-state Trp fluorescence emission spectra of 6 μM AstaPo1(apo), AstaPo1(ZEA) or AstaPo1^41-190^ (ZEA) were recorded upon excitation at 297 nm at 20 °C in the range of 305-450 nm at the rate of 30 nm/min on a Cary Eclipse spectrofluorometer (Varian) in the 20 mM HEPES buffer (pH 7.7) containing 150 mM NaCl. The spectrum of the buffer was subtracted from the protein spectra before presentation. The excitation and emission slits width was 5 nm. Indicated AstaPo1 concentrations are protein concentrations corrected for ZEA absorbance at 280 nm.

### 4.7. Differential scanning calorimetry

The AstaPo1 apoform (1.73 mg/ml, 80 μM) or its CAN-(1.66 mg/ml, 77 μM) or ZEA-bound (1.83 mg/ml, 85 μM) forms, or the truncated AstaPo1^41-190^ (ZEA) complex (1.3 mg/ml, 82 μM) were dialyzed overnight against a 20 mM Na-phosphate buffer (pH 7.0) and subjected to DSC on a VP-capillary DSC (Malvern) at a heating rate of 1 °C/min. Thermograms were processed using Origin Pro 8.0 and transition temperature (*T*_m_) was determined from the maximum of the thermal transition.

### 4.8. Small-angle X-ray scattering

SAXS data (I(s) versus s, where s = 4πsinθ/λ, 2θ is the scattering angle and λ = 1 Å) from AstaPo1(CAN) (2.54 mg/mL) or AstaPo1(ZEA) (7.05 mg/mL) were measured at 20 °C at the BM29 beamline (ESRF, Grenoble, France) using a Pilatus 2M detector in a batch mode. The buffer included 20 mM Tris-HCl, pH 7.6, and 150 mM NaCl. Series of frames for each sample revealed no significant radiation damage. Six statistically matching frames for each protein sample were averaged and buffer subtracted to produce the SAXS profile for further analysis of the SAXS-derived structural parameters, which are listed in **Table 2**. The SAXS data were processed and analyzed in PRIMUS (45). CRYSOL (46) was used for calculating theoretical SAXS profiles, fitting, and validation. To assess the quality and uniqueness of the approximation of the SAXS data by NMR structure models, alternative models based on the previously determined crystal structures of other carotenoproteins were used.

### 4.9. NMR spectroscopy

NMR spectra were recorded using the Bruker Avance III 600 MHz and 800 MHz spectrometers, both equipped with triple-resonance cryogenic probes. Spectra were acquired at two temperatures: 25 and 40 °C. Backbone NMR chemical shift assignment of AstaPo1, in both apo and holo forms, was performed using the conventional triple resonance approach (47), signals of aliphatic side chains were assigned based on 3D hCCH- and HcCH-TOCSY experiments and aromatic residues were assigned using the HCCH-COSY (48), (HB)CB(CGCCCar)Har (49) and 3D ^1^H,^15^N-NOESY-HSQC spectra. All the triple resonance spectra were acquired using the BEST-TROSY pulse sequences (50), and most of the 3D NMR spectra were recorded in a non-uniformly sampled regime and were processed using the qMDD software (51). For AstaPo1 in the apo state, the assignment of the protein in complex with AXT was used as an additional source of data. ^3^J_CCo_ and ^3^J_NCo_ couplings were measured from the spin-echo difference constant-time HSQC spectra (52, 53).^3^J_NHβ_ couplings were derived from the cross-peak intensities in a J-quantitative 3D HNHB experiment (54). To assign the chemical shift of AXT, we used the ^13^C,^15^N-double-filtered 2D NOESY, and TOCSY experiments (22). Intramolecular distances were measured in 3D ^1^H,^15^N-NOESY-HSQC, and ^1^H,^13^C-NOESY-HSQC. Intermolecular contacts were observed directly using the 3D ^13^C,^15^N-filtered,^13^C-edited-NOESY-HSQC (55).

Spatial structures were calculated using the automated procedure as implemented in CYANA version 3.98.13 (56). The intermolecular NOESY peak list was assigned manually. The dihedral restraints (*φ*, □_1_) were obtained from the manual analysis of J-couplings and chemical shifts in TALOS-N software (57). CYANA format library for the ligand was based on a library for the structure of AXT in complex with β-crustacyanin (25). This library was precalculated by the CYLIB algorithm (26). Values of dihedral angles were set according to a quantum chemical study (23). Dihedral angles of the ligand were restrained in the range (177;183) for most of the double bonds, the angle C5-C6-C7-C8 was restrained in the range (−173;-167) for the s-trans conformation and (−43;-37 or +37;+43) for the s-cis conformation. MOLMOL (58) and PyMOL software (Schrödinger LLC) was used for 3D visualization.

^15^N longitudinal (T1), and transverse (T2) relaxation rates of the AstaPo1/AXT complex were measured using the pseudo-3D HSQC-based experiments with varied relaxation delays at 25 °C using the Bruker Avance III 800 MHz spectrometer (59). Heteronuclear equilibrium ^1^H,^15^N-NOE magnitudes were obtained using the ^1^H presaturation for 3s during the recycling delay. Reference and NOE spectra were recorded in the interleaved mode. Relaxation parameters were analyzed using the model-free approach as implemented in the TENSOR2 software, assuming the isotropic rotation (60).

## Abbreviations used

βCar: β-carotene
BmCBP: *Bombyx mori* Carotenoid-Binding Protein
BME: β-mercaptoethanol
AstaP: astaxanthin-binding protein
AXT: astaxanthin
CAM: cell adhesion molecule
CAN: canthaxanthin
CD: circular dichroism
CTDH: C-terminal domain homolog
DMSO: dimethyl sulfoxide
ECN: echinenone
IMAC: immobilized metalaffinity chromatography
IPTG: isopropyl-β-thiogalactoside
MSA: multiple sequence alignment
MWCO: molecular weight cutoff
NDH-1: NAD(P)H dehydrogenase type 1
NMR: nuclear magnetic resonance
NOESY: Nuclear Overhauser Effect spectroscopy
OCP: Orange Carotenoid Protein
PDB: protein data bank
PMSF: phenylmethylsulfonyl fluoride
RMSD: root-mean-square deviation
SASA: solvent accessible surface area
SAXS: smallangle X-ray scattering
SDS-PAGE: sodium dodecyl sulfate-polyacrylamide gel electrophoresis
SEC: size-exclusion chromatography
TOCSY: total correlation spectroscopy
UV: ultraviolet
ZEA: zeaxanthin

## Acknowledgments

The authors thank Prof. Shinji Kawasaki for inspiration in studying AstaP and Dr. Anton Popov for help with SAXS data collection at the BM29 beamline (ESRF, Grenoble, France; experiment session data DOI 10.15151/ESRF-ES-496408264). The study was supported by the Ministry of Science and Higher education of the Russian Federation in the framework of the Agreement no. 075-15-2021-1354 (07.10.2021). CD measurements were done at the Shared-Access Equipment Centre “Industrial Biotechnology’’ of the Federal Research Center “Fundamentals of Biotechnology” of the Russian Academy of Sciences. SYK was supported by the Program of the Ministry of Science and Higher Education of Russia (0088-2021-0009). NMR studies were supported by the Grants Council of the President of the Russian Federation (grant M□-2834.2022.1.4).

## Data availability

Spatial structure of AstaPo1(AXT) complex and NMR chemical shifts were deposited to the Protein Data Bank under the access code 8C18 and to BMRB database under the access code 34781.

## Author contributions

Fedor D. Kornilov determined the structure and prepared figures

Yury B. Slonimskiy performed cloning and biochemical experiments, analyzed data

Daria A. Lunegova performed cloning and biochemical experiments

Nikita A. Egorkin performed cloning and discussed the results

Anna G. Savitskaya performed protein expression and purification

Sergey Yu. Kleymenov performed DSC

Eugene G. Maksimov obtained AXT, discussed the results

Sergey A. Goncharuk performed protein expression and purification, and supervision

Konstantin S. Mineev conceived studies, analyzed NMR data, wrote the paper, acquired funding and supervised research

Nikolai N. Sluchanko conceived studies, analyzed SAXS and biochemical data, wrote the paper, prepared figures, acquired funding and supervised research

## Supplementary data

### Supplementary Tables

**Table S1.**
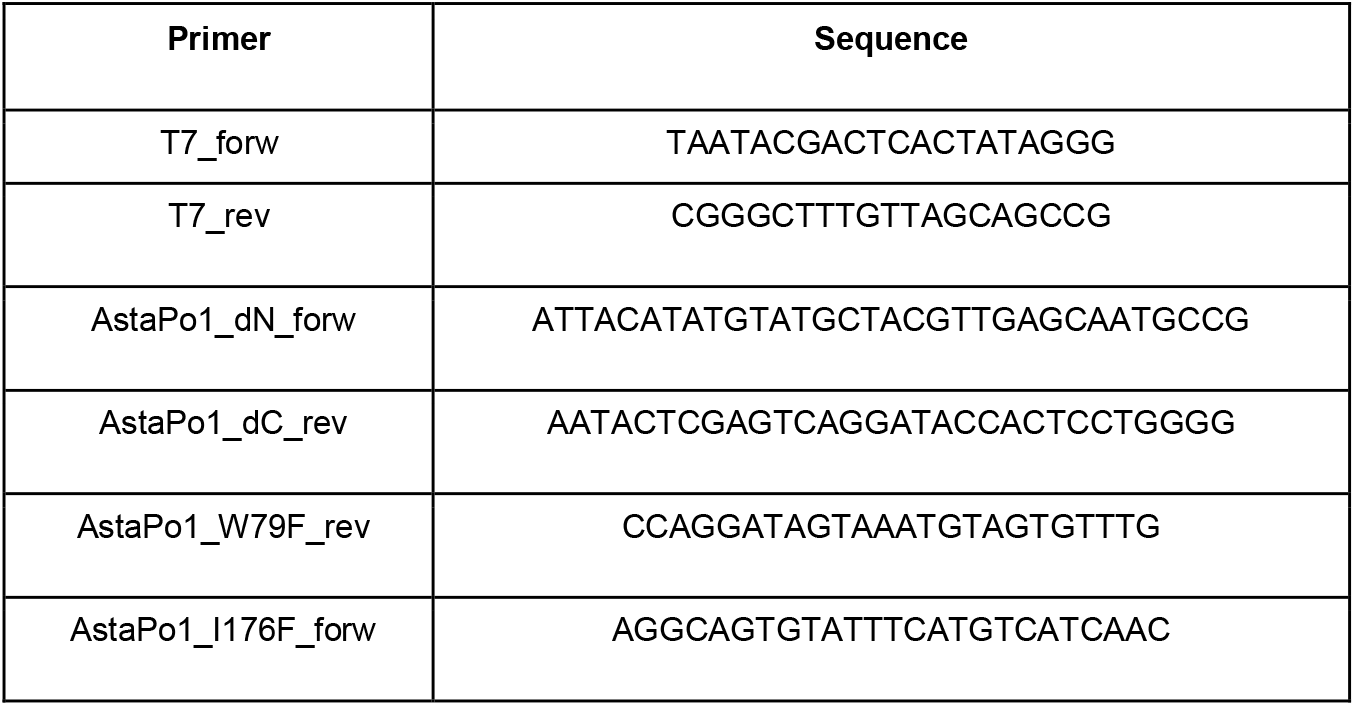
Primers used in this study.

**Table S2.** Properties of the proteins used in this study.

| Protein | $\epsilon_{280 \text{ nm}}$, $\text{M}^{-1} \text{cm}^{-1}$ | ZEA $A_{\text{vis}}/A_{\text{UV}}$ * | $M_w$ (kDa) | pI |
| --- | --- | --- | --- | --- |
| AstaPo1.WT | 22460 | 2.8 | 21.6 | 10.1 |
| AstaPo1.W79F | 16960 | ~3 | 21.6 | 10.1 |
| AstaPo1.I176F | 22460 | 2.8 | 21.6 | 10.1 |
| AstaPo1<sup>41-223</sup> | 22460 | 2.8 | 19.7 | 10.0 |
| AstaPo1<sup>21-190</sup> | 11460 | ~4 | 17.8 | 10.0 |
| AstaPo1<sup>41-190</sup> | 11460 | ~4 | 15.9 | 9.9 |
| SynAstaP (P74615) | 1490 | - | 16.3 | 4.7 |

All constructs contained GPHM extra residues at their N terminus, which was taken into account in calculations.
\*The expected Vis/UV absorbance ratios in ZEA-bound holoforms are indicated assuming the extinction coefficient of AstaPo1-bound ZEA at 463 nm and 280 nm equal to $120,000 \text{ M}^{-1} \text{cm}^{-1}$ and $20,000 \text{ M}^{-1} \text{cm}^{-1}$ , respectively, and taking into account protein extinction coefficients at 280 nm.

## Supplementary figures

**Fig. S1.**
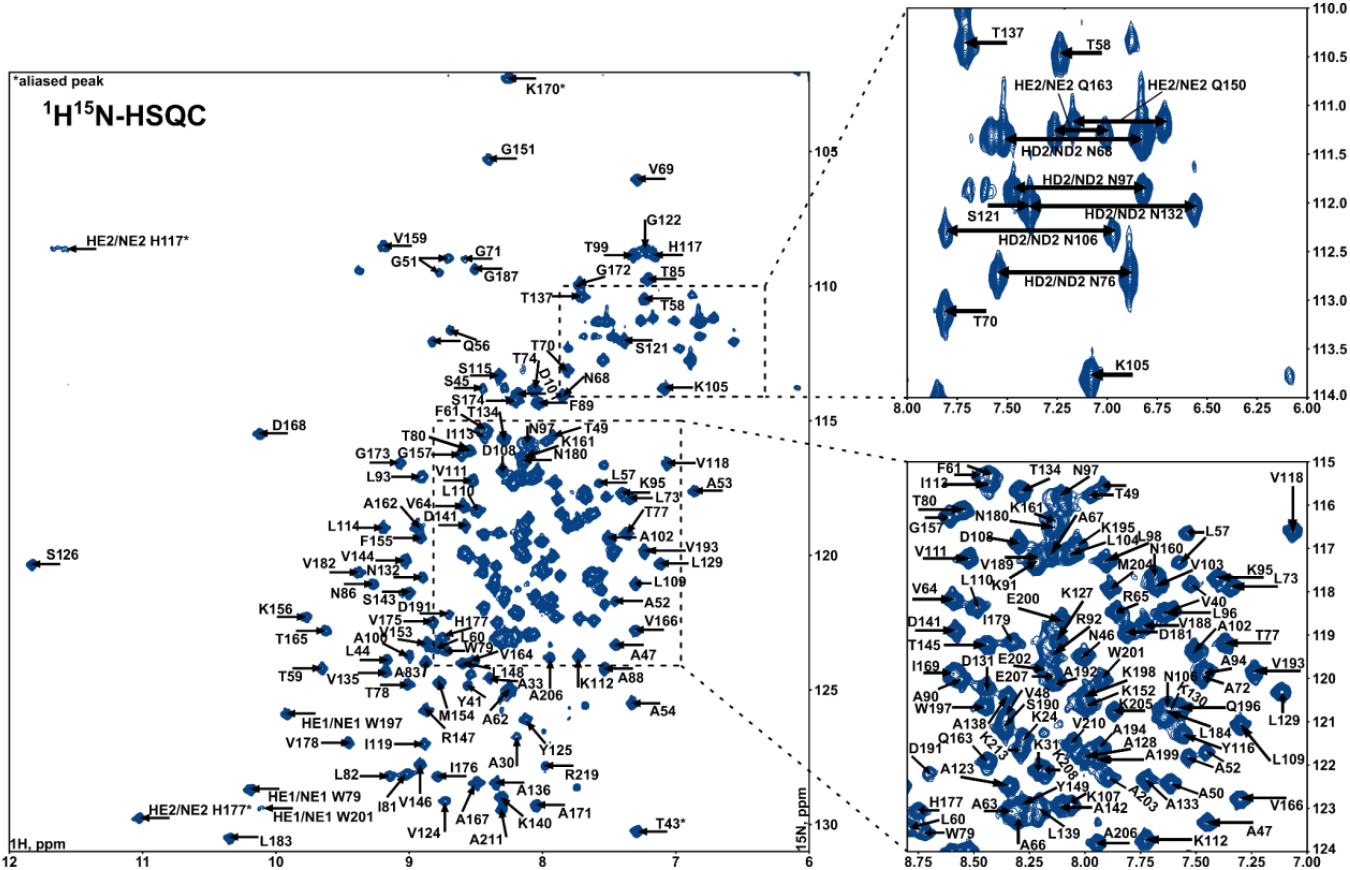
^1^H^15^N-HSQC spectrum of AstaPo1/AXT complex. T=313K, pH=7.5. Positions of aliased peaks of the histidine side chains are highlighted with asterisks.

**Supplementary text 1. Analysis of conformational heterogeneity of AXT in complex with AstaPo1.**

To elucidate the source of the conformational heterogeneity observed for the AstaPo1-bound AXT molecule, we considered three possible options: 1) the possibility that AstaPo1 can bind AXT in two alternative orientations, 2) the *s-cis-s-trans* isomerism of the C6-C7 bond and 3) the R-S optical isomers of astaxanthin, which are observed at the position 3 of the β-ionone group (**Fig. S2**). Inspection of the intermolecular distances suggests that the orientation of the isoprenoid fragment of AXT is the same in the two observed states, while the conformations or configurations of both β-ionone rings are different. To distinguish between the two remaining options, we analyzed the intra- and intermolecular distances and chemical shifts in AXT. Such an analysis reveals that one of the states of AXT corresponds to the (3S,3S)-AXT with the s-*cis* configuration of C6-C7 bonds for both rings. The s-cis conformation of AXT is more favorable (23) and corresponds to the state of the molecule in the DMSO solution (**Fig. S4**). The conformation of AXT in the second state is much less obvious. First of all, at one side of AXT, which contacts the helix a1, the conformation of the C6-C7 bond is s-cis in both states (the double bond proton, closest to the C17/C16 methyl groups is proximal to the isoprene methyl group). At the same time, the pattern of intermolecular NOEs observed for the H3-proton and C17/C16 methyl groups does not agree with the s-cis configuration of the bond and 3S-configuration of this β-ionone ring (**Fig. S5**). Thus, the only obvious option that agrees with all the data is the presence of a 3R-isomer of AXT (**Fig. S5**). According to the literature, both synthetic and natural AXT exists as a mixture of 3S and 3R isomers in various proportions (24), however, the exact isomer contents for the AXT preparation that we used in our research are not known. Apparently, in our particular case, the isomers are present in equal populations for both the headgroups of AXT. According to the ^13^Ca chemical shifts (**Fig. S6**), that are indicative of the protein secondary structure, AstaP adapts to the headgroup configuration by the subtle motions (the chemical shift difference is as small as 0.3 ppm) in the hinge region connecting helices a1 and a2.

**Fig. S2.**
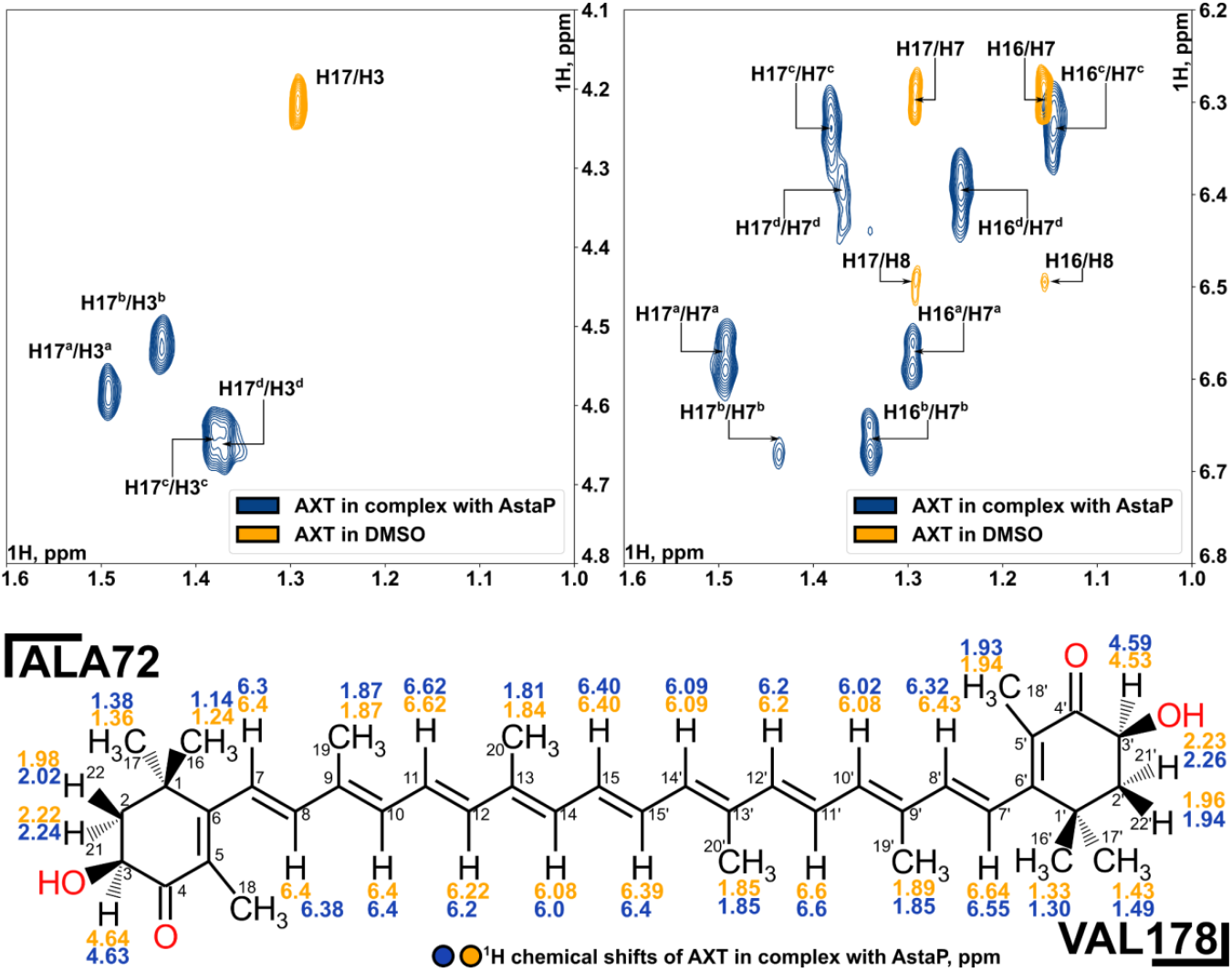
Fragments of the 2D ^13^C/^15^N-double filtered-NOESY spectra recorded for the AstaPo1/AXT complex in water (shown in blue) and of the 2D-NOESY spectra recorded for the AXT in DMSO (shown in yellow). Four sets of signals corresponding to AXT in complex with AstaPo1 are indicated by the superscripts a-d. The structure of AXT with the atom numbering is also shown. Assignment of the proton chemical shift for the two states of AXT is shown by yellow and blue numbers, Ala72 and Val178 denote the residues that are in contact with the corresponding AXT headgroup in both states, according to the ^13^C-filtered NOESY experiment.

**Fig. S3.**
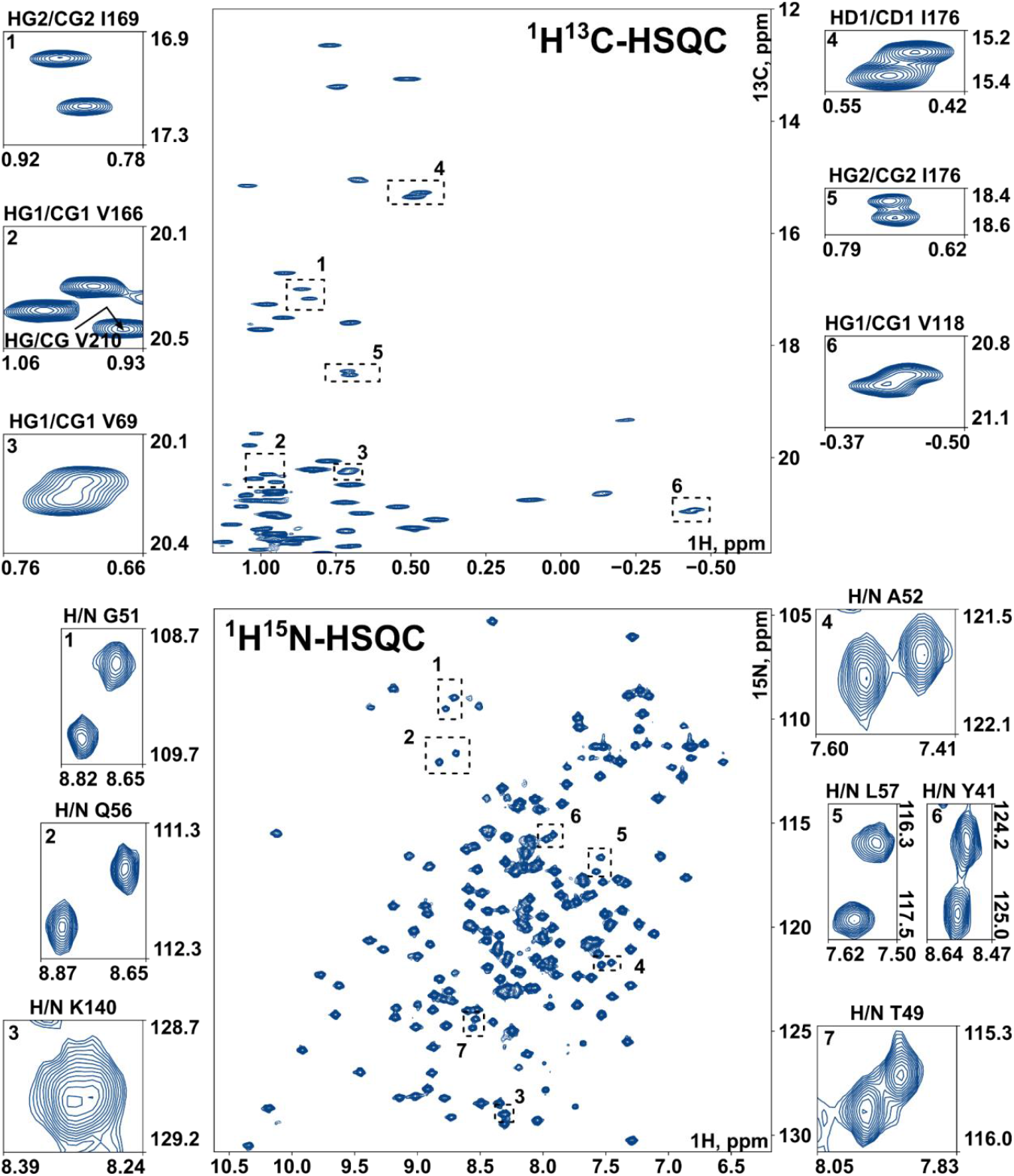
Signal splitting in ^1^H,^13^C-CT-HSQC (a fragment containing the signals from the Leu, Ile, Val methyl groups) and ^1^H,^15^N-HSQC spectra of AstaPo1/AXT complex recorded at 40° C. Split signals are shown by numbered dashed rectangles and are additionally shown at high resolution.

**Fig. S4.**
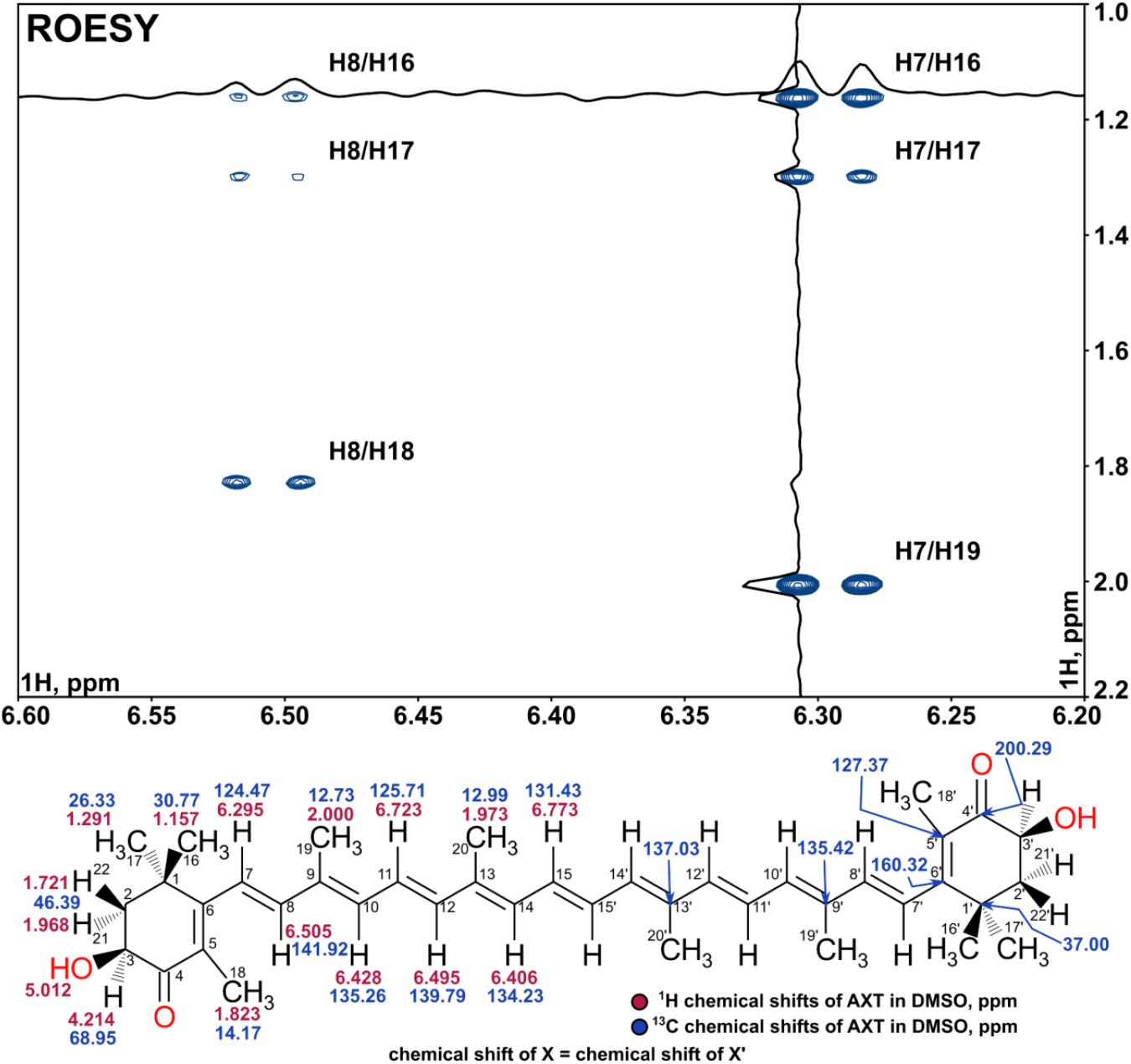
A fragment of ROESY spectrum of 1 mM AXT solution in DMSO, showing the contacts between the C1 (H16 and H17), C5 (H18), and C9 (H19) methyl groups in the AXT headgroup and H7/H8 protons of the double bond. The analysis supports the s-cis configuration of the C6-C7 bond. The structure of AXT and chemical shift assignments are shown below.

**Fig. S5.**
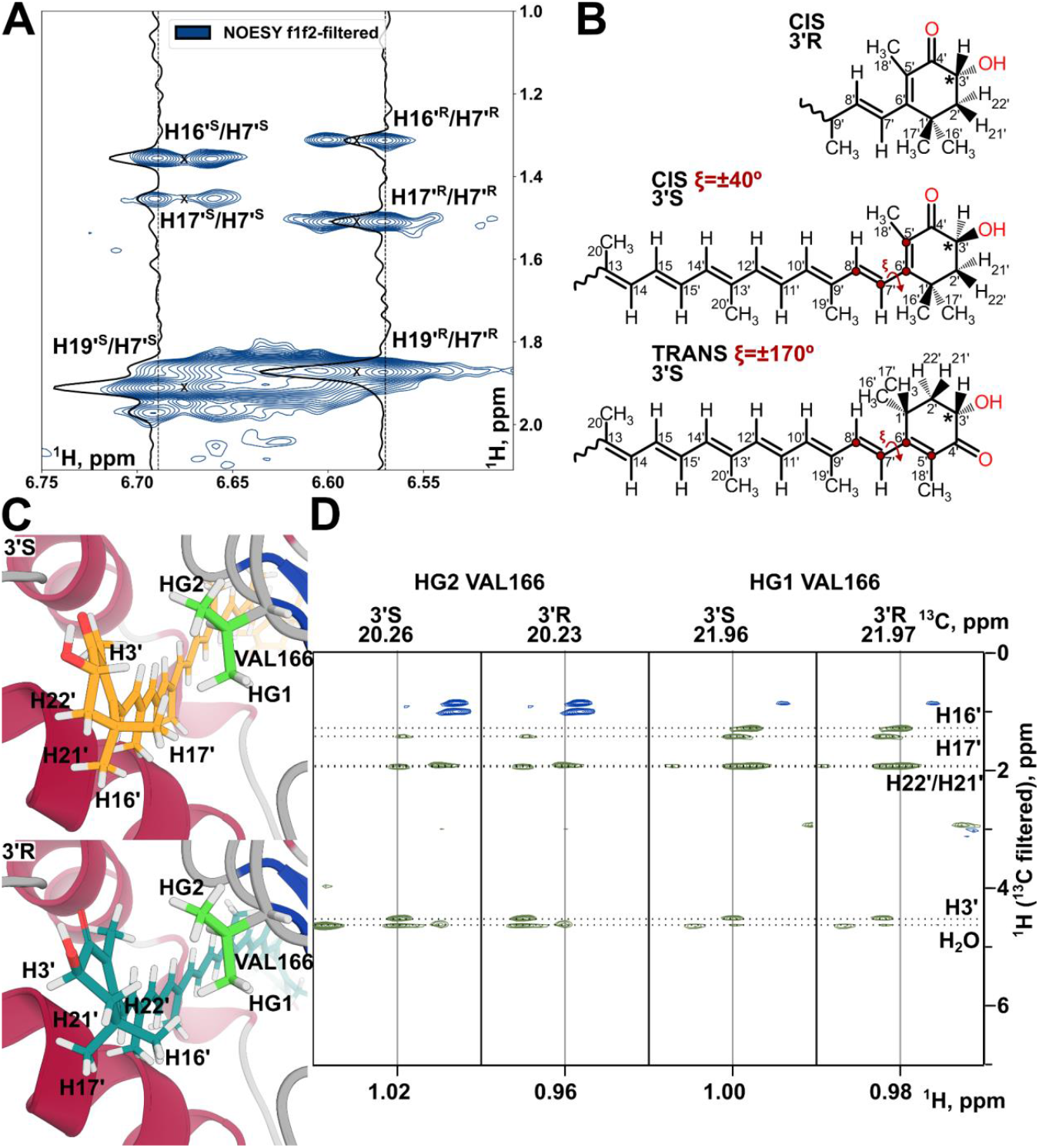
Analysis of AXT configurations in complex with AstaPo1. **A** - a fragment of 2D f1f2-filtered NOESY spectrum of AstaPo1/AXT complex, showing the NOE-contacts between H7’ and methyl groups (H16’, H17’ and H19’) of AXT. The strongest peak is observed with the H19 methyl group, supporting the *cis* conformation of C6-C7 bond. **B** - Cis/Trans conformations and 3’S/3’R isomers of AXT. The stereocenter is marked with an asterisk, red circles indicate atoms that define a dihedral angle ξ. **C** - 3’S (from the NMR structure) and 3’R (putative model) stereoisomers of AXT in the exit from the binding pocket of AstaPo1 contact the Val166 sidechain. **D** - 2D 1H/1H strips from the 3D 13C/15N-filtered,13C-edited-NOESY-HSQC, showing intermolecular NOE-contacts between HG1/HG2 of VAL166 and atoms of the β-ionone ring of AXT. Contacts to the H3’ proton are observed only in the 3’S state.

**Fig. S6.**
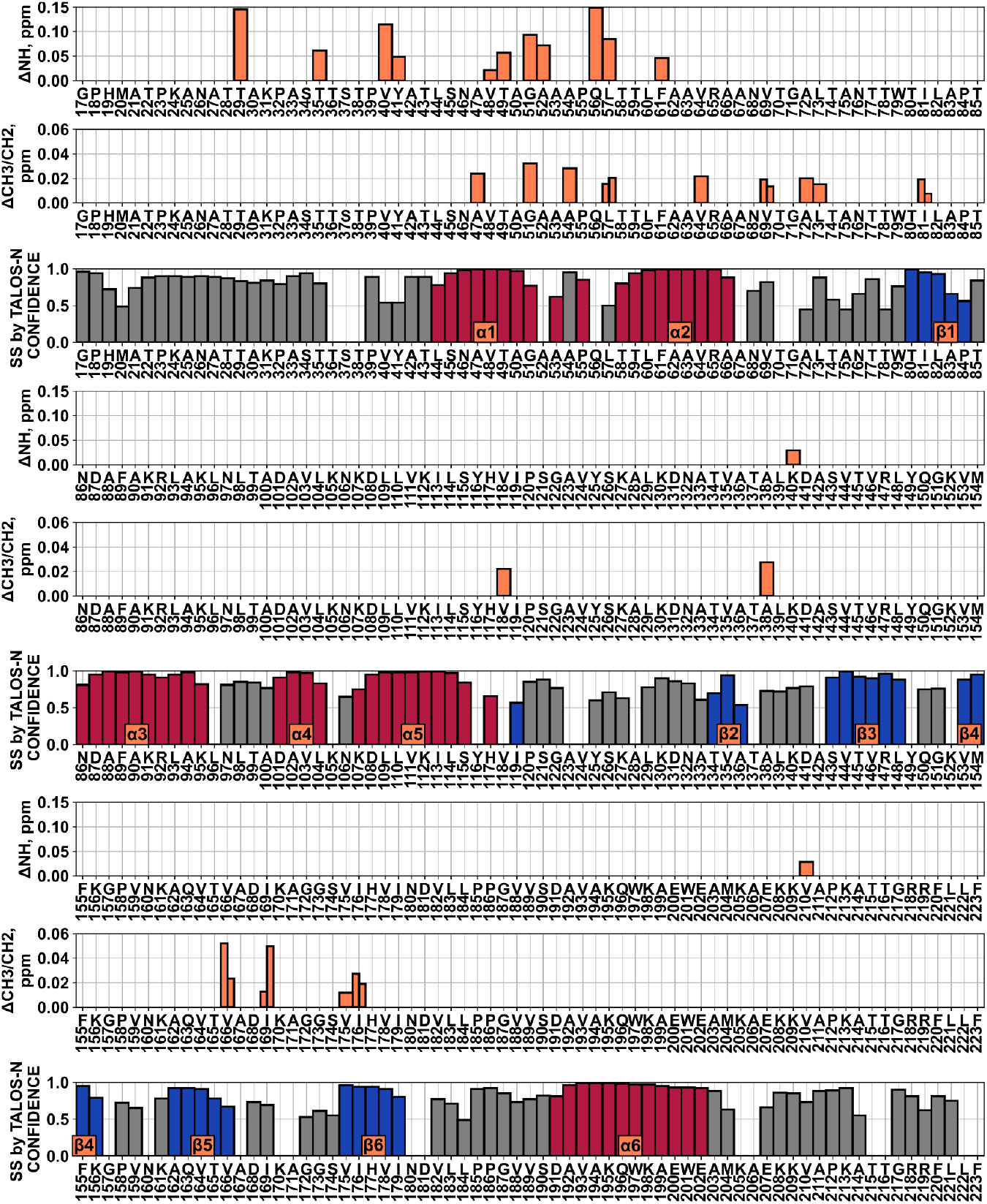
Peak splittings in the NMR spectra of AstaPo1/AXT complex. Chemical shift perturbations of AstaPo1 amide and methyl groups that take place due to the conformational heterogeneity of AstaPo1/AXT complex. Additionally, the secondary structure is provided, as predicted by TALOS-N software based on the NMR chemical shifts. Bar height corresponds to the confidence value and bar color indicates the predicted type of spatial structure.

**Fig. S7.**
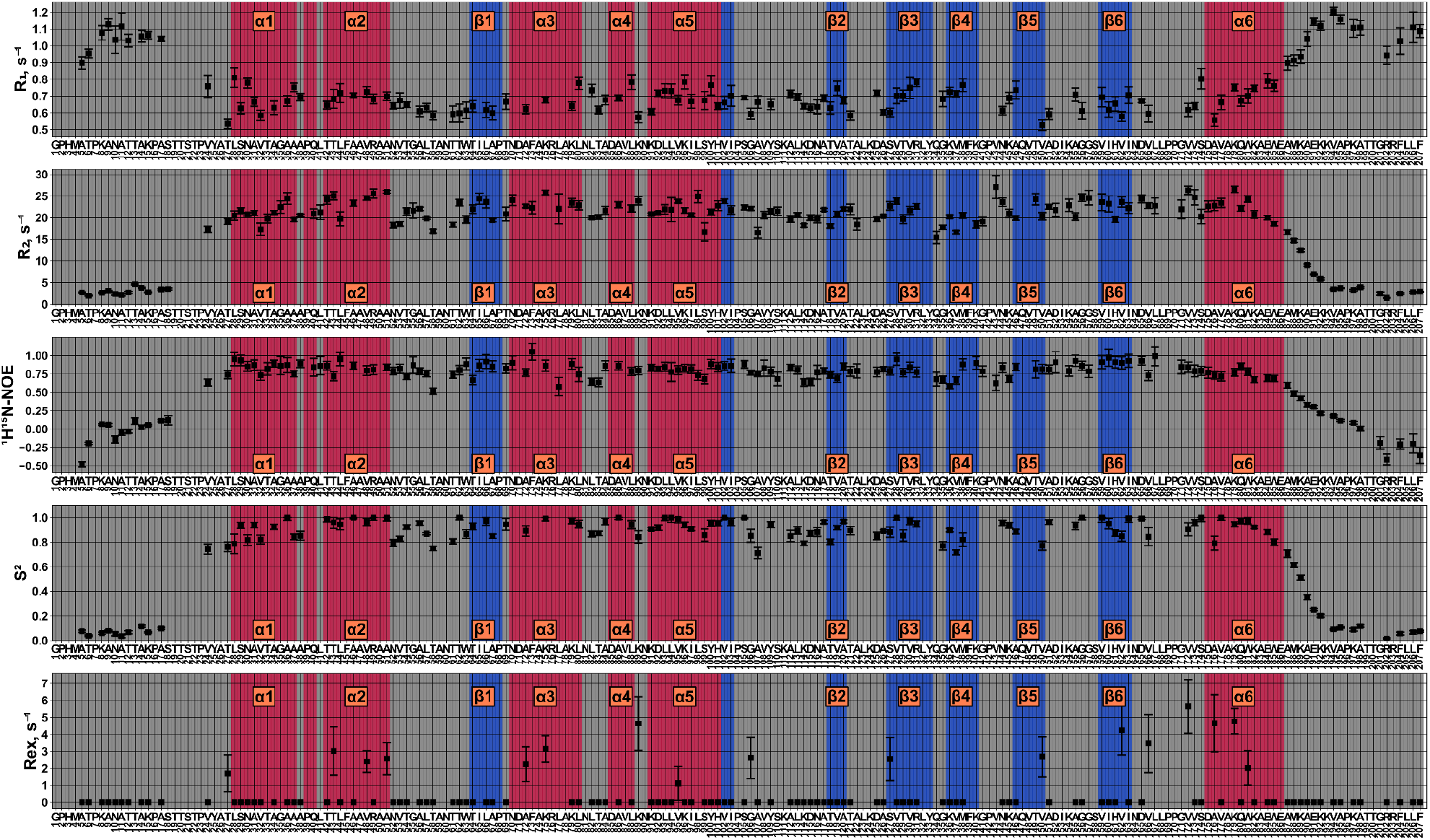
Per-residue dynamics of the AstaPo1/AXT complex. NMR relaxation parameters of ^15^N nuclei (rates of longitudinal (R1) and transverse (R2) relaxation, heteronuclear equilibrium NOE (^1^H,^15^N NOE)) and internal mobility parameters - generalized order parameter S^2^ and exchange contribution to the transverse relaxation R_ex_ measured for AstaPo1 in complex with AXT. The elements of the secondary structure are indicated by red (helices) and blue (strands) bars.

**Fig. S8.**
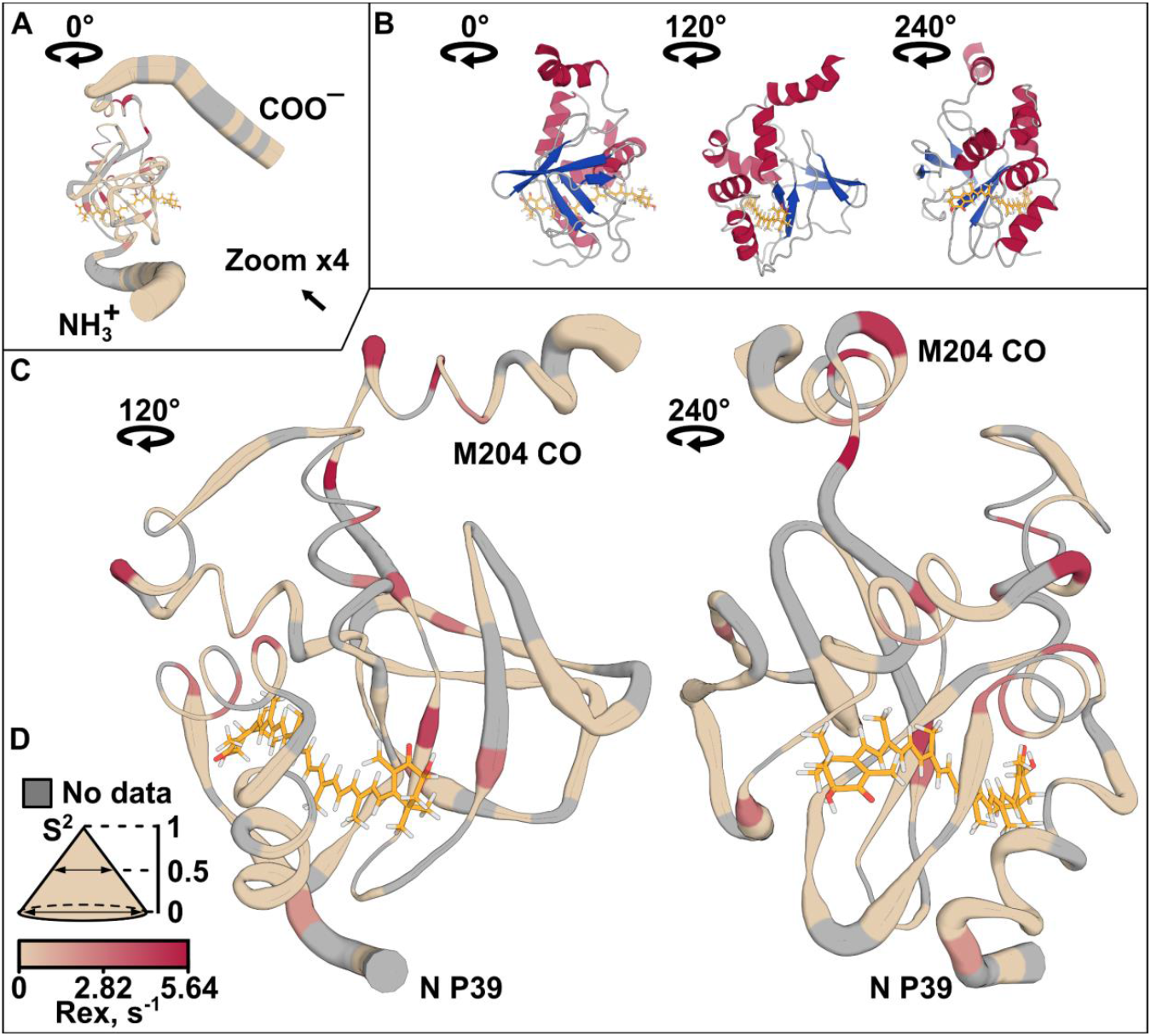
Dynamics of the AstaPo1(AXT) structure. **A,C** - “sausage plot”, highlighting the dynamics of the AstaPo1/AXT backbone. The thickness of the ribbons is inversely proportional to the generalized order parameters (S^2^) of HN bonds. Residues with significant contributions of slow motions to the transverse relaxation (Rex) are colored red, according to the displayed scale. Grey regions correspond to the residues with no data available (prolines, overlapped, and too broad signals). Mobile terminal regions are removed on panel **C** for clarity. Panel **B** displays the cartoon of the views shown in **A** and **C**, denoted as 0, 120 and 240°.

**Fig. S9.**
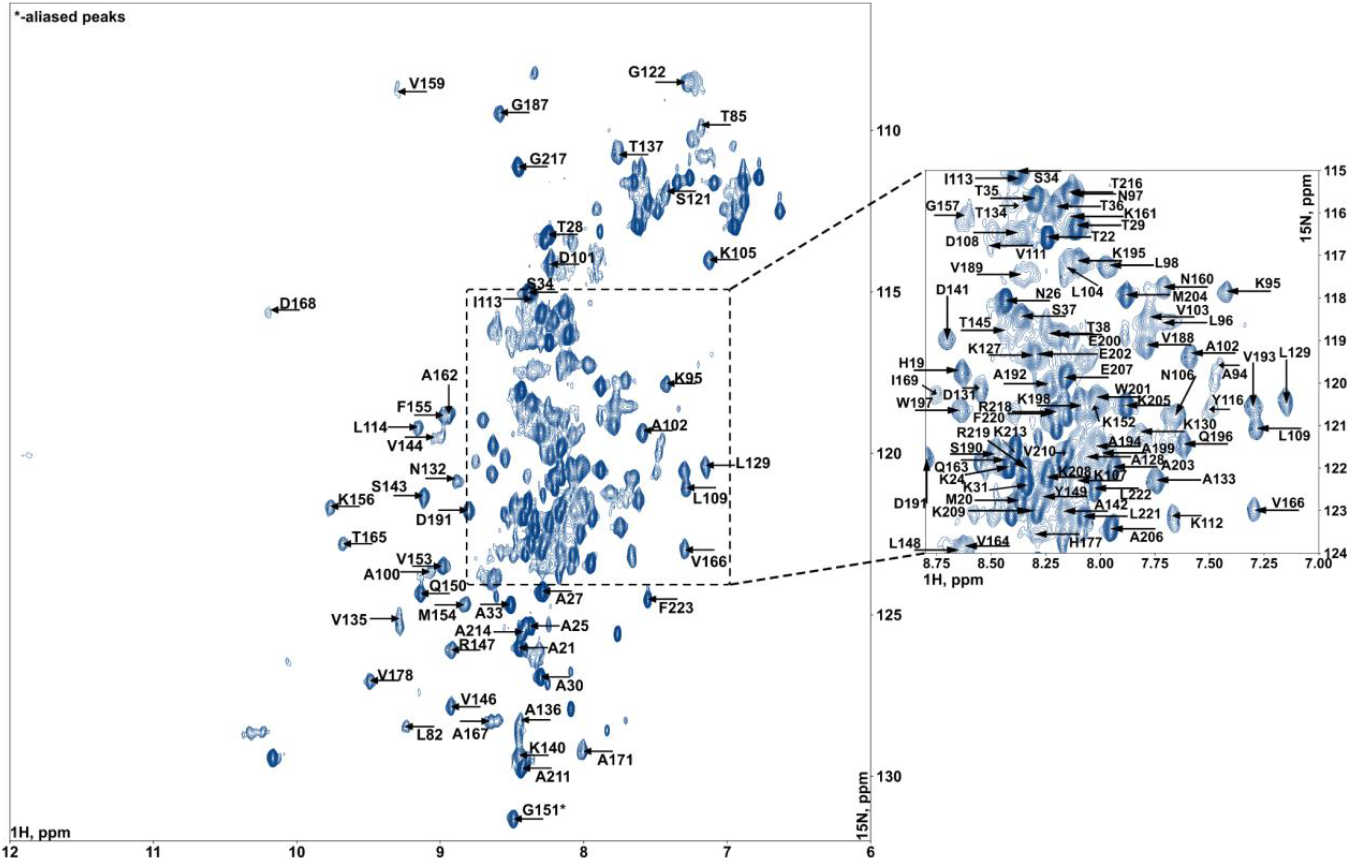
^1^H^15^N-HSQC spectrum of AstaPo1(apo). T=298K, pH=6.0. Positions of aliased peaks are highlighted with asterisks.

**Fig. S10.**
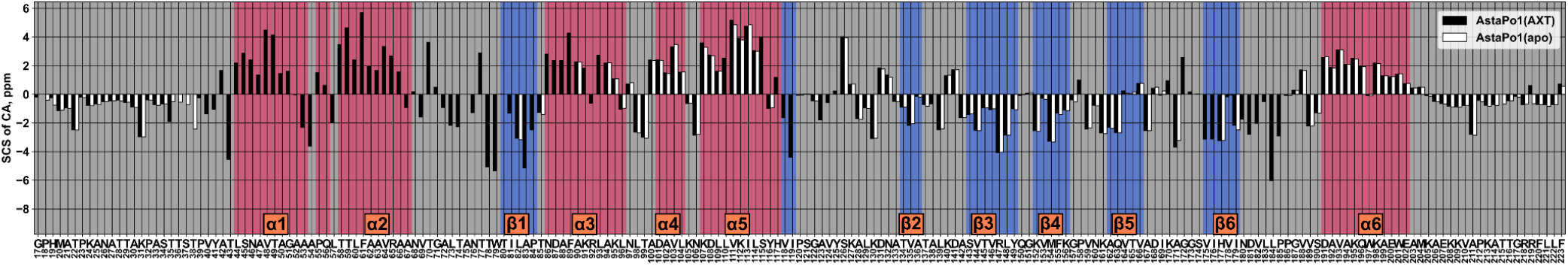
Comparison of ^13^Ca secondary chemical shifts (SCS) between AstaPo1(AXT) (black color) and AstaPo1(apo) (white color).

**Fig. S11.**
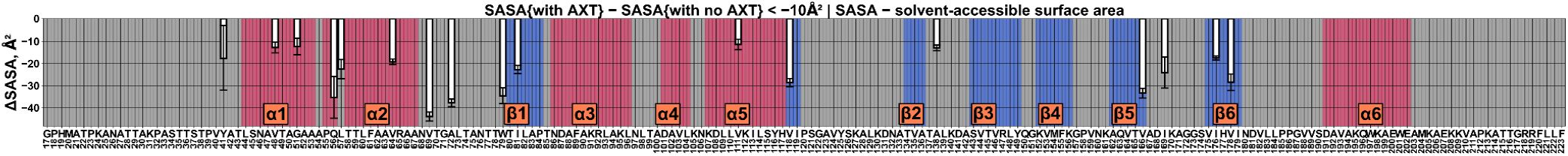
Solvent-accessible surface area (SASA) changes with and without carotenoid using 10 A^2^ as cutoff).

## References

1. M. J. Bastiani, A. L. Harrelson, P. M. Snow, C. S. Goodman, Expression of fasciclin I and II glycoproteins on subsets of axon pathways during neuronal development in the grasshopper. Cell 48, 745–755 (1987).

2. G. J. Seifert, Fascinating Fasciclins: A Surprisingly Widespread Family of Proteins that Mediate Interactions between the Cell Exterior and the Cell Surface. Int. J. Mol. Sci. 19, 1628 (2018).

3. N. J. Clout, D. Tisi, E. Hohenester, Novel Fold Revealed by the Structure of a FAS1 Domain Pair from the Insect Cell Adhesion Molecule Fasciclin I. Structure 11, 197–203 (2003).

4. O. Politz, et al., Stabilin-1 and −2 constitute a novel family of fasciclin-like hyaluronan receptor homologues. Biochem. J. 362, 155–164 (2002).

5. M. D. Carr, et al., Solution Structure of the Mycobacterium tuberculosis Complex Protein MPB70: FROM TUBERCULOSIS PATHOGENESIS TO INHERITED HUMAN CORNEAL DISEASE *. J. Biol. Chem. 278, 43736–43743 (2003).

6. R. G. Moody, M. P. Williamson, Structure and function of a bacterial Fasciclin I Domain Protein elucidates function of related cell adhesion proteins such as TGFBIp and periostin. FEBS Open Bio 3, 71–7 (2013).

7. A. Korste, H. Wulfhorst, T. Ikegami, M. M. Nowaczyk, R. Stoll, Solution structure of the NDH-1 complex subunit CupS from Thermosynechococcus elongatus. Biochim. Biophys. Acta BBA - Bioenerg. 1847, 1212–1219 (2015).

8. R. García-Castellanos, et al., Structural and Functional Implications of Human Transforming Growth Factor β-Induced Protein, TGFBIp, in Corneal Dystrophies. Structure 25, 1740–1750.e2 (2017).

9. J. Liu, et al., Structural characterizations of human periostin dimerization and cysteinylation. FEBS Lett. 592, 1789–1803 (2018).

10. O. Huber, M. Sumper, Algal-CAMs: isoforms of a cell adhesion molecule in embryos of the alga Volvox with homology to Drosophila fasciclin I. EMBO J. 13, 4212–4222 (1994).

11. A. Faik, J. Abouzouhair, F. Sarhan, Putative fasciclin-like arabinogalactan-proteins (FLA) in wheat (Triticum aestivum) and rice (Oryza sativa): identification and bioinformatic analyses. Mol. Genet. Genomics 276, 478–494 (2006).

12. S.-Y. Park, M.-Y. Jung, I.-S. Kim, Stabilin-2 mediates homophilic cell–cell interactions via its FAS1 domains. FEBS Lett. 583, 1375–1380 (2009).

13. S. Kawasaki, K. Mizuguchi, M. Sato, T. Kono, H. Shimizu, A novel astaxanthin-binding photo-oxidative stress-inducible aqueous carotenoprotein from a eukaryotic microalga isolated from asphalt in midsummer. Plant Cell Physiol 54, 1027–40 (2013).

14. S. Kawasaki, et al., Photooxidative stress-inducible orange and pink water-soluble astaxanthin-binding proteins in eukaryotic microalga. Commun Biol 3, 490 (2020).

15. H. Toyoshima, et al., Distribution of the Water-Soluble Astaxanthin Binding Carotenoprotein (AstaP) in Scenedesmaceae. Mar Drugs 19 (2021).

16. H. Toyoshima, S. Takaichi, S. Kawasaki, Water-soluble astaxanthin-binding protein (AstaP) from Coelastrella astaxanthina Ki-4 (Scenedesmaceae) involving in photooxidative stress tolerance. Algal Res. 50, 101988 (2020).

17. Y. B. Slonimskiy, N. A. Egorkin, T. Friedrich, E. G. Maksimov, N. N. Sluchanko, Microalgal protein AstaP is a potent carotenoid solubilizer and delivery module with a broad carotenoid binding repertoire. FEBS J. 289, 999–1022 (2022).

18. E. A. Burstein, N. S. Vedenkina, M. N. Ivkova, Fluorescence and the location of tryptophan residues in protein molecules. Photochem Photobiol 18, 263–79 (1973).

19. C. A. Kerfeld, et al., The crystal structure of a cyanobacterial water-soluble carotenoid binding protein. Structure 11, 55–65 (2003).

20. D. Harris, et al., Structural rearrangements in the C-terminal domain homolog of Orange Carotenoid Protein are crucial for carotenoid transfer. Commun Biol 1, 125 (2018).

21. N. N. Sluchanko, et al., Structural basis for the carotenoid binding and transport function of a START domain. Structure 30, 1647–1659.e4 (2022).

22. J. Iwahara, J. M. Wojciak, R. T. Clubb, Improved NMR spectra of a protein–DNA complex through rational mutagenesis and the application of a sensitivity optimized isotope-filtered NOESY experiment. J. Biomol. NMR 19, 231–241 (2001).

23. B. Durbeej, L. A. Eriksson, Protein-bound chromophores astaxanthin and phytochromobilin: excited state quantum chemical studies. Phys. Chem. Chem. Phys. 8, 4053–4071 (2006).

24. V. M. Moretti, et al., Determination of astaxanthin stereoisomers and colour attributes in flesh of rainbow trout (Oncorhynchus mykiss) as a tool to distinguish the dietary pigmentation source. Food Addit. Contam. 23, 1056–1063 (2006).

25. M. Cianci, et al., The molecular basis of the coloration mechanism in lobster shell: beta-crustacyanin at 3.2-A resolution. Proc Natl Acad Sci U A 99, 9795–800 (2002).

26. E. M. Yilmaz, P. Güntert, NMR structure calculation for all small molecule ligands and non-standard residues from the PDB Chemical Component Dictionary. J. Biomol. NMR 63, 21–37 (2015).

27. N. N. Sluchanko, et al., OCP-FRP protein complex topologies suggest a mechanism for controlling high light tolerance in cyanobacteria. Nat Commun 9, 3869 (2018).

28. M. Moldenhauer, et al., Assembly of photoactive orange carotenoid protein from its domains unravels a carotenoid shuttle mechanism. Photosynth Res 133, 327–341 (2017).

29. H. Ashkenazy, et al., ConSurf 2016: an improved methodology to estimate and visualize evolutionary conservation in macromolecules. Nucleic Acids Res 44, W344–50 (2016).

30. E. D. Drew, R. W. Janes, PDBMD2CD: providing predicted protein circular dichroism spectra from multiple molecular dynamics-generated protein structures. Nucleic Acids Res. 48, W17–W24 (2020).

31. M. R. Melnicki, et al., Structure, Diversity, and Evolution of a New Family of Soluble Carotenoid-Binding Proteins in Cyanobacteria. Mol Plant 9, 1379–1394 (2016).

32. F. Muzzopappa, et al., Paralogs of the C-Terminal Domain of the Cyanobacterial Orange Carotenoid Protein Are Carotenoid Donors to Helical Carotenoid Proteins. Plant Physiol 175, 1283–1303 (2017).

33. Y. B. Slonimskiy, et al., Light-controlled carotenoid transfer between water-soluble proteins related to cyanobacterial photoprotection. FEBS J 286, 1908–1924 (2019).

34. E. G. Maksimov, et al., NMR resonance assignment and backbone dynamics of a C-terminal domain homolog of orange carotenoid protein. Biomol. NMR Assign. 15, 17–23 (2021).

35. M. A. Dominguez-Martin, et al., Structural analysis of a new carotenoid-binding protein: the C-terminal domain homolog of the OCP. Sci Rep 10, 15564 (2020).

36. E. G. Maksimov, et al., Soluble Cyanobacterial Carotenoprotein as a Robust Antioxidant Nanocarrier and Delivery Module. Antioxid. Basel 9 (2020).

37. M. A. Dominguez-Martin, et al., Structural and spectroscopic characterization of HCP2. Biochim Biophys Acta Bioenerg 1860, 414–424 (2019).

38. T. Sakudoh, et al., Carotenoid silk coloration is controlled by a carotenoid-binding protein, a product of the Yellow blood gene. Proc. Natl. Acad. Sci. 104, 8941–8946 (2007).

39. F. Alpy, C. Tomasetto, Give lipids a START: the StAR-related lipid transfer (START) domain in mammals. J Cell Sci 118, 2791–801 (2005).

40. K. V. Tugaeva, N. N. Sluchanko, Steroidogenic Acute Regulatory Protein: Structure, Functioning, and Regulation. Biochem. Mosc 84, S233–S253 (2019).

41. N. N. Sluchanko, et al., Silkworm carotenoprotein as an efficient carotenoid extractor, solubilizer and transporter. Int. J. Biol. Macromol. 223, 1381–1393 (2022).

42. P. Régnier, et al., Astaxanthin from Haematococcus pluvialis Prevents Oxidative Stress on Human Endothelial Cells without Toxicity. Mar. Drugs 13, 2857–2874 (2015).

43. L.-Y. Zang, O. Sommerburg, F. J. G. M. van Kuijk, Absorbance Changes of Carotenoids in Different Solvents. Free Radic. Biol. Med. 23, 1086–1089 (1997).

44. F. W. Studier, Protein production by auto-induction in high density shaking cultures. Protein Expr Purif 41, 207–34 (2005).

45. D. Franke, et al., ATSAS 2.8: a comprehensive data analysis suite for small-angle scattering from macromolecular solutions. J Appl Crystallogr 50, 1212–1225 (2017).

46. D. I. Svergun, C. Barberato, M. H. J. Koch, CRYSOL - a Program to Evaluate X-ray Solution Scattering of Biological Macromolecules from Atomic Coordinates. J Appl Cryst 28, 768–773 (1995).

47. L. E. Kay, M. Ikura, R. Tschudin, A. Bax, Three-dimensional triple-resonance NMR spectroscopy of isotopically enriched proteins. J. Magn. Reson. 1969 89, 496–514 (1990).

48. K. Pervushin, R. Riek, G. Wider, K. Wüthrich, Transverse Relaxation-Optimized Spectroscopy (TROSY) for NMR Studies of Aromatic Spin Systems in 13C-Labeled Proteins. J. Am. Chem. Soc. 120, 6394–6400 (1998).

49. F. Löhr, R. Hänsel, V. V. Rogov, V. Dötsch, Improved pulse sequences for sequence specific assignment of aromatic proton resonances in proteins. J. Biomol. NMR 37, 205–224 (2007).

50. A. Favier, B. Brutscher, Recovering lost magnetization: polarization enhancement in biomolecular NMR. J. Biomol. NMR 49, 9–15 (2011).

51. M. Mayzel, K. Kazimierczuk, V. Y. Orekhov, The causality principle in the reconstruction of sparse NMR spectra. Chem. Commun. 50, 8947–8950 (2014).

52. G. W. Vuister, A. C. Wang, A. Bax, Measurement of three-bond nitrogen-carbon J couplings in proteins uniformly enriched in nitrogen-15 and carbon-13. J. Am. Chem. Soc. 115, 5334–5335 (1993).

53. S. Grzesiek, G. W. Vuister, A. Bax, A simple and sensitive experiment for measurement of JCC couplings between backbone carbonyl and methyl carbons in isotopically enriched proteins. J. Biomol. NMR 3, 487–493 (1993).

54. P. Düx, B. Whitehead, R. Boelens, R. Kaptein, G. W. Vuister, Measurement of 15N-1H coupling constants in uniformly 15N-labeled proteins: Application to the photoactive yellow protein. J. Biomol. NMR 10, 301–306 (1997).

55. C. Zwahlen, et al., Methods for Measurement of Intermolecular NOEs by Multinuclear NMR Spectroscopy: Application to a Bacteriophage λ N-Peptide/boxB RNA Complex. J. Am. Chem. Soc. 119, 6711–6721 (1997).

56. P. Güntert, L. Buchner, Combined automated NOE assignment and structure calculation with CYANA. J. Biomol. NMR 62, 453–471 (2015).

57. Y. Shen, A. Bax, “Protein Structural Information Derived from NMR Chemical Shift with the Neural Network Program TALOS-N” in Artificial Neural Networks, Methods in Molecular Biology., H. Cartwright, Ed. (Springer, 2015), pp. 17–32.

58. R. Koradi, M. Billeter, K. Wüthrich, MOLMOL: A program for display and analysis of macromolecular structures. J. Mol. Graph. 14, 51–55 (1996).

59. N. A. Farrow, et al., Backbone Dynamics of a Free and a Phosphopeptide-Complexed Src Homology 2 Domain Studied by 15N NMR Relaxation. Biochemistry 33, 5984–6003 (1994).

60. P. Dosset, J.-C. Hus, M. Blackledge, D. Marion, Efficient analysis of macromolecular rotational diffusion from heteronuclear relaxation data. J. Biomol. NMR 16, 23–28 (2000).

